# Essential role of Bone morphogenetic protein 15 in porcine ovarian and follicular development and ovulation

**DOI:** 10.1101/724096

**Authors:** Yufeng Qin, Tao Tang, Wei Li, Zhiguo Liu, Xiaoliang Yang, Xuan Shi, Guanjie Sun, Xiaofeng Liu, Min Wang, Xinyu Liang, Peiqing Cong, Delin Mo, Xiaohong Liu, Yaosheng Chen, Zuyong He

## Abstract

Bone morphogenetic protein 15 (BMP15) is a multifunctional oocyte-specific secreted factor. It controls female fertility and follicular development in both species-specific and dosage-sensitive manners. Previous studies found that BMP15 played a critical role on follicular development and ovulation rate of mono-ovulatory mammalian species, but has minimal impact on poly-ovulatory mice. However, whether this is true in non-rodent poly-ovulatory species need to be validated. To investigate this question, we generated a BMP15 knockdown pig model. We found that BMP15 knockdown gilts showed markedly reduced fertility accompanied with phenotype of dysplastic ovaries containing significantly declined number of follicles, increased number of abnormal follicles, and abnormally enlarged antral follicles resulting in disordered ovulation. Molecular and transcriptome analysis revealed that knockdown of *BMP15* significantly suppressed cell proliferation, differentiation, *Fshr* expression, leading to premature luteinization and reduced estradiol production in GCs, and simultaneously decreased the quality and meiotic maturation of oocyte. Our results provide *in vivo* evidences for the essential role of BMP15 in porcine ovarian and follicular development, and new insight into the complicated regulatory function of BMP15 in female fertility of poly-ovulatory species.

## Introduction

In the past three decades, increasing studies have revealed the important role of the oocyte-specific secreted factor BMP15 in mammalian ovarian and follicular development through exerting its multiple functions including promoting granulosa cells proliferation and steroidogenesis(Moore et al., 2003; Moore and Shimasaki, 2005; Otsuka et al., 2001; Otsuka et al., 2000), preventing cell apoptosis and premature luteinization(Chang et al., 2013; Hussein et al., 2005; Juengel et al., 2011; McNatty et al., 2005; Zhai et al., 2013), regulating glycometabolism and lipid metabolism(Su et al., 2008; Sugiura et al., 2007), controlling oocyte competence and ovulation(Fabre et al., 2006a; Hussein et al., 2006). As a key signaling molecular mediating the dialogue between oocyte and its surrounding somatic cells(Gilchrist et al., 2008), BMP15 expresses initially in early follicle stage, and gradually increases in subsequent follicle stages to the period of ovulation and/or luteinization(Paradis et al., 2009; Sun et al., 2010). This expression pattern is a little different in species, for example, initial expression of BMP15 protein can be found in primary follicle stage of sheep, human and pig, but didn’t in mice until pre-ovulatory stage(Paulini and Melo, 2011). BMP15 protein secretes and functions as BMP15/BMP15 homodimers and BMP15/GDF9 (growth differentiation factor 9) heterodimers, through binding to the membrane bound type II serine/threonine kinase BMP receptor (BMPR2) and type I activin receptor-like kinase ALK6, resulting in the phosphorylation and activation of SMAD pathways(Liao et al., 2003; Mottershead et al., 2013; Pulkki et al., 2012). In particular, BMP15 homodimers are considered to bind to ALK6 receptor to activate the Smad1/5/8 signaling pathway in some species, for example in human and sheep but not in rodent. While BMP15/GDF9 heterodimers are considered to bind to BMPR2 receptor to activate Smad2/3 signaling pathway in all reported species (sheep, human, mouse, pig at al)(Lin et al., 2014; Peng et al., 2013; Reader et al., 2011; Reader et al., 2016). In most cases, BMP15/GDF9 heterodimers were more potent in regulation of GCs, oocyte, and zygote development(Peng et al., 2013).

BMP15 mutations or deficience has been associated with altered female fertility in different species. As previously reported, natural mutations in BMP15 of sheep can lead to increased ovulation rate and litter size in heterozygotes, but infertility in homozygotes due to bilateral ovarian hypoplasia(Braw-Tal et al., 1993; Fabre et al., 2006b; Galloway et al., 2000; Smith et al., 1997). Altered fertility also has been reported in sheep immunized with BMP15 mature protein or different region of peptides(Juengel et al., 2002; Juengel et al., 2004; Juengel et al., 2013; McNatty et al., 2007). In human, BMP15 mutations have been associated with primary ovarian insufficiency (POI) and infertility phenotype of women(Abir et al., 2014; Al-ajoury et al., 2015; Chand et al., 2006). However, in the poly-ovulatory mice, there was no significant difference between *BMP15^+/−^* females and wild-type in ovulation rate, and only a mild reduction of fertility in *BMP15* null female mice(Yan et al., 2001). Several studies have attempted to determine whether there are species-specific differences in the BMP15 system that may play causal roles in the differences in fertility observed in mono-ovulatory mouse and poly-ovulatory sheep and humans. One study has attributed the species-specific differences to the temporal variations in the production of the mature form of BMP15. They found that mouse BMP15 mature protein was barely detectable until preovulatory stage, when it is markedly increased(Yoshino et al., 2006). They subsequently found that defects in the production of mouse BMP15 mature protein could correlate with species-specific differences(Hashimoto et al., 2005). Moreover, a phylogenetic analysis found that a better conservation in areas involved in dimer formation and stability of BMP15 within mono-ovulatory species, but high variations in these areas within poly-ovulatory species, implying the correlation with altered equilibrium between homodimers and heterodimers, and modified biological activity for allowing polyovulation to occur(Monestier et al., 2014). Hence, it seems that the role of BMP15 in regulation of follicular development and ovulation rate was more critical in mono-ovulatory mammalian species than poly-ovulatory animals. However, whether BMP15 is essential to ovarian and follicular development in poly-ovulatory mammalian species still remains unclear, as this has not yet been tested in *in vivo* studies of non-rodent poly-ovulatory mammals.

In this study, we aim to investigate the function of BMP15 on female fertility and follicular development of non-rodent poly-ovulatory mammal by using a *BMP15* knockdown transgenic pig model. The transgenic (TG) gilts appeared decreased female fertility with phenotypes of disordered estrous cycle, significant reduced ovarian size and follicle number, higher ratio of abnormal follicles, and none corpus lutein formed before 365 days old. We found that knocking down of *BMP15* can impair porcine follicle growth and cause dysovulation mainly by affecting oocyte quality and oocyte meiotic maturation, and suppressing GCs proliferation and GCs functions, including inhibiting the expression of *Fshr* and E2 production, resulting in premature luteinization. These effects on follicular cell functions could finally lead to absence of dominant follicle selection but appearance of abnormally enlarged antral follicles with ovulation dysfunction in transgenic gilts. Our findings were evidently different from the unchanged fertility of *BMP15^+/−^* mice, strongly suggesting the important role of BMP15 in non-rodent poly-ovulatory mammals, thus providing the basis for further investigation of the different regulatory role of BMP15 between mono-and poly-ovulatory mammals.

## RESULTS

### Generation and identification of *BMP15* knockdown pig model

To generated the *BMP15* RNA interference transgenic pig model, we first designed and constructed 5 pEGFP-*BMP15-*shRNA plasmids, in which *BMP15* shRNA sequence was under control of human U6 promoter, then inserted downstream of the *EGFP* expression cassette (Fig. 1A). Each shRNA expression plasmids was respectively cotransfected into HEK293T cell with a psiCheck II-*BMP15* plasmid to examine their RNA interference efficiency *in vitro*. We found shRNA1 was most effective with a RNA inference efficiency reaching to 76% (Fig. 1B), thus this shRNA was selected for transfection into embryonic fibroblast cells (PEFs) derived from a male Yorkshire pig. Transfected PEFs then were subjected to G418 selection to screen the cells with stable expression of EGFP as donor cells for somatic cell nuclear transfer (SCNT). Clone embryos then were transferred into Large White sow recipients to generate F0 TG pigs as described in our previous report(Liu et al., 2019) (Fig. S1A). We obtained two healthy F0 TG males at last. After sexual maturity, one F0 TG boar was mated with wild-type sows to generate F1 TG gilts for subsequent experiments. Both F0 and F1 TG pigs showed visible intense GFP fluorescence on toes and muscle while subjected to sunlight (Fig. 1C, Fig. S1B), directly suggesting that the pEGFP-*BMP15* shRNA plasmid was successfully integrated into the genome of F0 TG boar, and can be transmitted to the next generation through the germline. This was confirmed by PCR analysis of fragment of integrated plasmid in muscle tissue of F1 TG gilts (Fig. S2A). The copy number of integrated plasmid was estimated to be approximate seven in F1 TG pigs through the combination of both qPCR that using a *transferrin receptor* gene to normalize the genomic DNA (data not shown), and Southern blot analysis (Fig. S2B). More importantly, evidently decrease level of *BMP15* mRNA (Fig. 1E) in 365 days old TG ovaries and BMP15 protein level in 30 days old TG ovaries (Fig. 1F, G) strongly demonstrated the successful generation of the *BMP15* knockdown model, and implied an *in vivo* BMP15 knockdown efficiency of about 50% in TGF ovaries. Our qPCR data also revealed that *BMP15* mRNA was highly expressed in ovary tissue, exhibited a very low level in the pituitary (Fig. 1D), but was undetectable in another porcine tissues (e.g., liver, muscle, kidney).

**Fig. 1.**
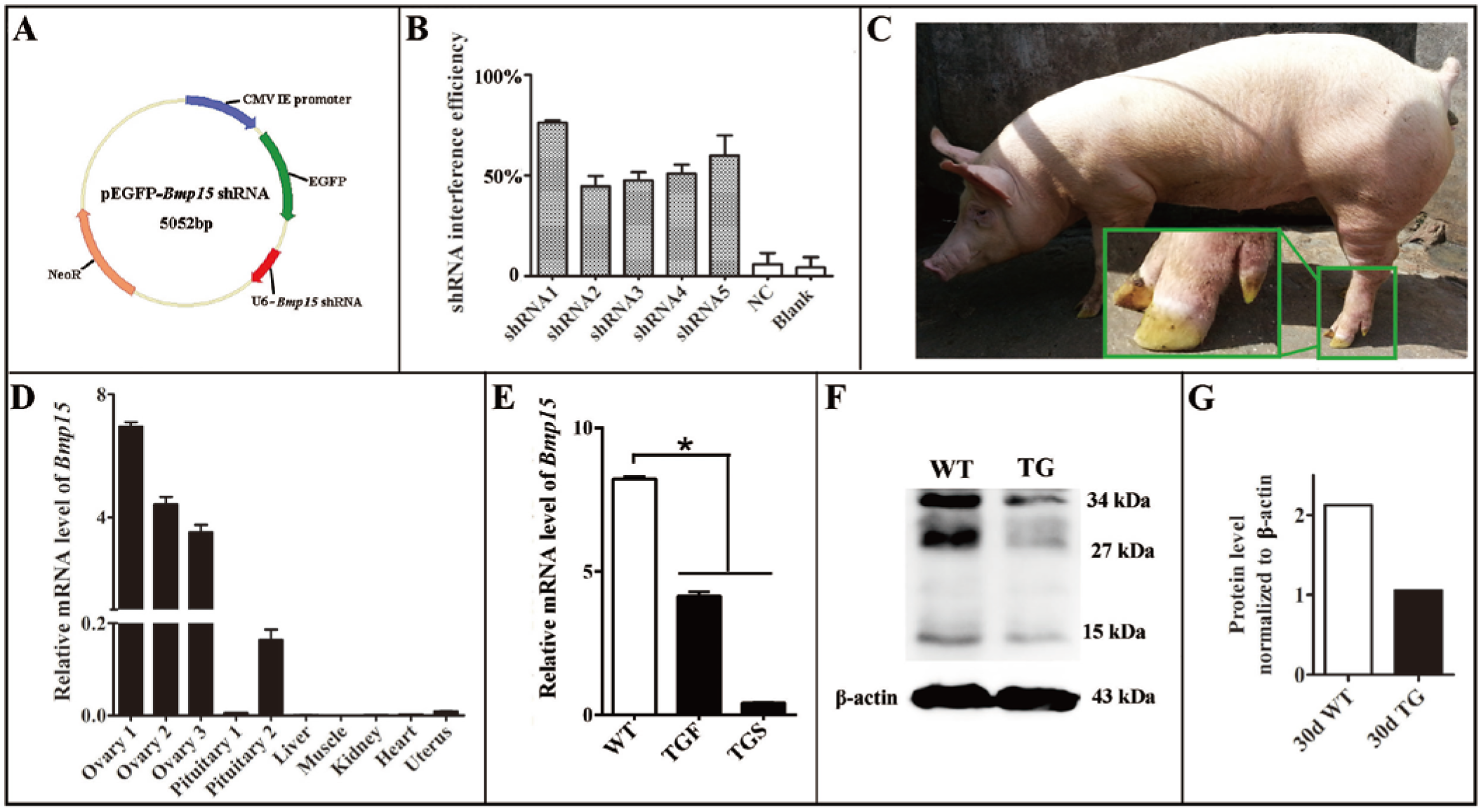
Generation of the *BMP15* knockdown pig model. **(A)** Diagram of shRNA expression vector. Synthesized *hU6*-*BMP15* shRNA fragment was inserted downstream of *EGFP* expression cassette on pEGFP-N1 vector. **(B)** RNA interference efficiency of 5 *BMP15* shRNAs was examined by a dual-luciferase reporter system after 48 h transfection of h293T cells. NC, random shRNA plasmid. **(C)** F1 TG gilt showed a visible GFP fluorescence on the toes while under sunlight. **(D)** Tissue-specific mRNA expression profile of BMP15 WT pigs. **(E)** qPCR analysis of *BMP15* mRNA level in 365 days old transgenic ovaries with two different phenotypes (TGF and TGS). TGF, transgenic ovary with many visible antral follicles on ovarian surface. TGS, transgenic ovary with streak phenotype. *P < 0.05. **(F)** Western blot analysis of BMP15 protein level in postnatal 30 days old TG ovaries. Three prominent, distinct bands were observed corresponding to apparent molecular weights of 34 kDa, 27 kDa, and 15 kDa. **(G)** Quantitative analysis of BMP15 protein levels based on the band intensity in f by using Image J software.

### Knockdown of *BMP15* was associated with disordered reproductive cycle of TG gilts

All F1 TG gilts presented normal appearance and growth condition, and 50 of them at age of 170 to 400 days old were checked daily for signs of oestrus in the presence of an intact mature boar. Surprisingly, we didn’t find any obvious estrous behavior or vulvar appearance changes (e.g., increased redness, swelling or mucus production) in sexually mature TG gilts (Fig. S3A). About twenty gilts at age of 240 to 400 days were bred by artificial insemination (AI) after treatment with PG600, but all failed to become pregnant. To determine whether the disordered estrous cycle in TG gilts was related to altering reproductive hormone changes, we measured the concentration of plasma estrogen (E2), progesterone (P4) and follicle stimulating hormone (FSH) in 365 days old gilts throughout the estrous cycle. The results showed that a typical peripheral E2 concentration peak before the onset of estrous can be observed in WT gilts, which is consistent with previous studies(Soede et al., 2011) (Fig. 2A). In contrast, irregular E2 concentration peaks were observed in TG gilts during the continuous 24 days measurement (Fig. 2B). Vaginal smears cytology analysis(Mayor et al., 2007) further proved the disordered estrous cycle occurred in TG gilts, as irregular cytologic changes was observed through 16 days continuous examination (Fig. S3C). Furthermore, the average level of peripheral P4 concentration was significant lower in two TG gilts, (Fig. 2C), while higher serum FSH concentration was found in two of the three TG gilts (Fig. S3B). In addition, we found over 2-fold up-regulated expression of *Fsh* mRNA level in the pituitary of both 150 and 260 day TG gilts (Fig. 2D), but no significant difference in the expression level of *luteinizing hormone* (*Lh*) simultaneous (Fig. 2E). These results indicated a disordered reproductive cycle and potential ovarian dysfunction in TG gilts.

**Fig. 2.**
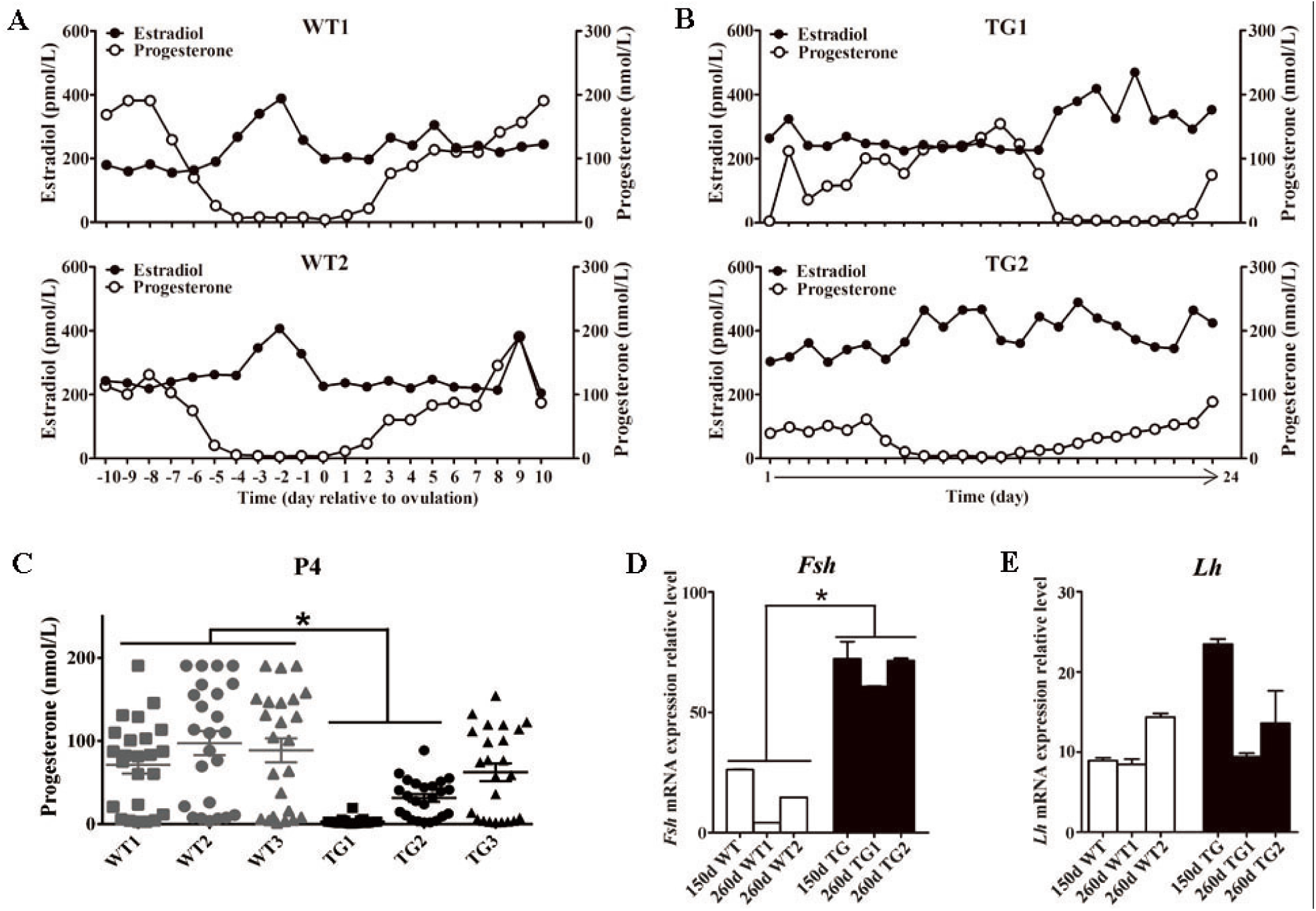
TG gilts presented disordered estrous cycle and reproductive hormones. Plasma E2 and P4 concentration of 365-day old WT and TG gilts were mesured at a 24h interval for 24 days. **(A)** During the estrous cycle, two representative WT gilts showed typical serum E2 concentration peak before ovulation, accompanied with marked decreased P4 concentration. **(B)** Irregular plasma E2 concentration peaks was observed in two representative TG gilts in continuous 24 days measurement. **(C)** The average P4 concentration of two of the three TG gilts in continuous 24 days measurement was significantly lower than WT gilts. (P<0.05). Each point stands for a P4 concentration value. **(D)** Expression level *o*f *Fsh* mRNA in the pituitary of both 150 and 260 day TG gilts were all more than 2-fold higher than WT gilts. (P<0.05). **(E)** The average level of *Lh* mRNA level in pituitary was not significantly different between TG and WT gilts.

### Knockdown of *BMP15* led to inhibition of follicular development and ovulation in TG ovaries

Since the estrous cycle is determined by ovarian and follicular development (Noguchi et al., 2010), the disordered estrous cycle of TG gilts potentially caused by impaired ovarian follicular development. In this regard, ovaries from gilts of different ages were collected and processed for morphological examination. Surprisingly, we found remarkably decreased size in TG ovarian and number of antral follicles (AFs) on the surface of TG ovaries of 140 to 365-day old gilts (Fig. 3A). In addition, apparent size difference was observed between bilateral TG ovaries (Fig. 3A). Corpus lutein was not observed in TG ovaries from 140 to 365 days old gilts, but can be found in 400 and 500 day TGF ovaries (Fig. S4A). Besides, the weight of TG ovaries before sexual maturity was markedly lower than WT ovaries (Table S4). Among the TG ovaries, we discovered 8 streak ovaries, denoted as TGS ovaries, in 6 gilts at age of 110 to 365 days, presenting an incidence of about 14%, while no streak ovary was found in WT ovaries (Fig. 3A,B). These TGS ovaries contained none or less than 3 visible AFs on the ovarian surface. In cortex of TGS ovaries from 110 and 200-day old gilts, most follicles were arrested in primary stage (Fig. 3B, E). In cortex of TGS ovaries from 365 day old gilts most follicles were arrested in secondary stage, and degradation of follicles became apparent (Fig. 3B). In different to TGS ovaries, the rest of TG ovaries contained many visible large AFs on the surface. We denoted them as TGF ovaries (Fig. 3A, C). Different stage of follicles can be found in these TGF ovaries, however, the follicle number decreased drastically during follicular development (Fig. 3D, E). Notably, during the early follicle stage, the significantly decline of primordial and primary follicle number led to a much thinner ovarian cortex in TGF ovaries of pre-puberty gilts. In addition, structural abnormality of SFs was evident, particularly in TGF ovaries of puberty gilts (Fig. 3C, G, H, Fig. S4B). We observed abnormally enlarged (Fig. 3H) or degenerated oocytes (Fig. 3Gii), multioocytic follicles (Fig. 3Gi) highly irregular GC layers (Fig. 3Gii,iv) and degraded GCs (Fig. 3Giv), and abnormally thickened theca layers (Fig. 3Giii). These follicular abnormalities in certain extent were similar to those found in previous studies on animals with natural BMP15 mutations and immunized with BMP15 peptides(Juengel et al., 2009; Juengel et al., 2002; McNatty et al., 2007; Smith et al., 1997). Furthermore, a statistical counting on the ovarian sections from 5 TG gilts at age of 160 to 400 days showed an markedly reduced proportion of normal SFs in the TGF ovaries (Fig. 3F).

**Fig. 3.**
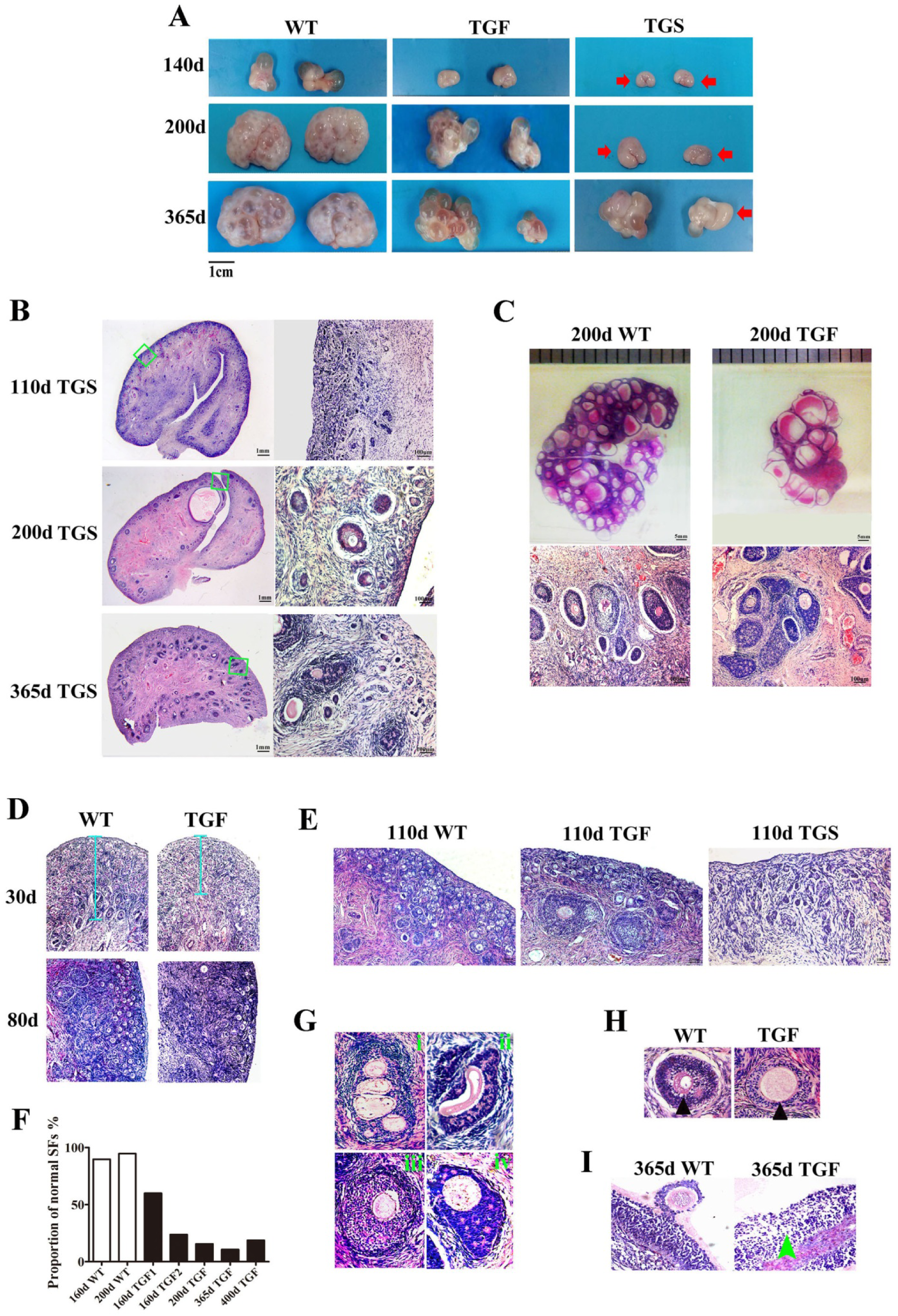
Affected ovarian and follicular development by knocking down of BMP15. **(A)** Representative photograms of ovaries collected from gilts of different ages showed reduced ovarian size and less visible follicles on the surface of TG ovaries as compared to ovaries from WT sibling. Bilateral TG ovaries were significantly different in size at age of 200 and 365 days. Two ovarian phenotypes were identified, TGF ovaries had many visible large antral follicles on the ovarian surface. TGS ovaries contained none or less than three visible antral follicles (red arrows). **(B)** Histological observation of TGS ovaries showed that 110-day TGS ovary presented major primary-like follicles sparsely scattered on the cortex, while 365-day TGS ovarian section was predominantly occupied by degraded secondary follicles. **(C)** On 200-day TGF ovarian section, decreased number of follicles, while enlarged antrum of antral follicles was observed. In addition, degradation of GCs in abnormally organized GC layer structure of secondary follicles was oberved. **(D)** In 30 and 80-day TGF ovaries, drastically decreased number of early stage follicles led to thinner ovarian cortex (blue line). **(E)** Comparison of three ovarian phenotypes at age of 110 days showed less number of early stage follicles in TGF ovarian cortex, and the minimum number of follicles in TGS ovaries. **(F)** Results of a follicle number counting showed markedly reduced proportion of normal secondary follicles in the TGF ovaries. Secondary follicles in three sections of each ovary were counted. **(G)** Representative images of abnormal TGF secondary follicles, including multiovular follicle with highly irregularly organized theca cell layers (i); follicle with oocyte-free structure, and abnormally thickened zona pellucida surrounded by highly degraded GCs (ii); follicle with abnormally thickened theca layers (iii); follicle with enlarged oocyte surrounded by highly irregularly organized GC layers with holes formed by degradation of GCs (iv). **(H)** TGF follicle showed larger oocyte in the early secondary follicle stage (black arrow head). **(I)** Smaller GCs were loosely organized in TGF antral follicles (green arrow).

Histological observation also revealed some striking features of TGF AFs. Most notably, AF number declined remarkably, but its antrum was enlarged substantially (Fig. 3C), and surrounded by loosely organized smaller GCs (Fig. 3I). We further isolated the AFs from three 365-day TGF ovaries derived from different gilts for statistical analysis. The results showed that TGF ovaries contained less total number of AFs, and also the number of small AFs (diameter<5mm), however, it contained substantially more large AFs with diameter >5mm (Fig. 4A, B). We found the diameter of the largest AF in TGF ovary can reach to 9 mm, while this was about 7 mm in WT ovaries (Fig. 4C). Normally, porcine AFs stop growth at diameter about 5mm, only selected follicles continue to grow through accumulation of follicular fluid, and ovulate at a diameter about 7mm(Soede et al., 2011). Thus the increased abnormally enlarged AFs in the TGF ovaries may be related to dysovulation and the disordered serum reproductive hormones found in TG gilts. Subsequent measurement of the concentration of reproductive hormones in follicular fluid of TGF large AFs (diameter >5 mm) showed that the E2 concentration was remarkably lower (Fig. 4D), but the concentration of other three hormones including P4 (Fig. 4E), FSH (Fig. S4C) and LH (Fig. S4D) all was not significant different from that in WT large AFs. Reduced E2 production in these TGF large follicles may imply an absence of dominant follicle selection(Clement and Monniaux, 2013). Taken these results together, we provided convincing evidence that knocking down of *BMP15* could severely inhibit both follicular development and ovulation of polyo-vulatory pig.

**Fig. 4.**
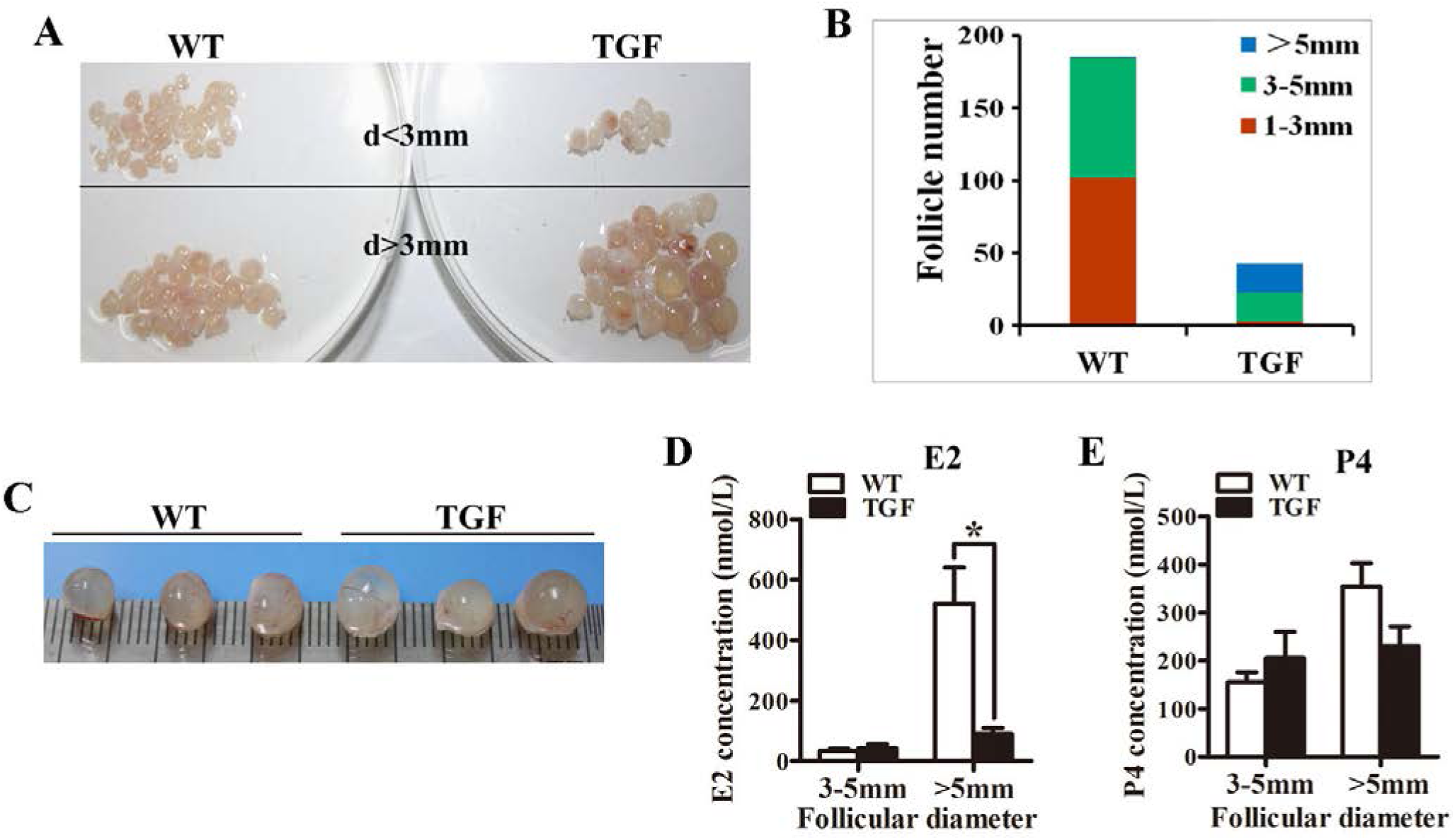
Transgenic ovaries contained abnormally enlarged antral follicles with dramatically reduced concentration of follicular fluid E2. **(A)** Photograms of the isolated antral follicle of 365-day ovaries demonstrated remarkably declined number of antral follicles with a diameter <3 mm in TGF ovaries. **(B)** Statistical data showed reduced total number of antral follicles, and substantially increased number of follicles with a diameter > 5mm in 365-day TGF ovaries. Antral follicles were isolated from three ovaries of different gilts, and then classified into 3 groups according to their diameter (d 1-3 mm, d 3-5mm, d>5mm). **(C)** Comparison of three largest follicles isolated from WT and TGF ovaries. **(D)** E2 concentration in follicular fluid of TGF large antral follicles was significantly lower than that in WT pre-ovulatory follicles. * stands for P<0.05. **(E)** P4 concentration in follicular fluid was not significantly different between TGF and WT. Hormones in follicular fluid were measured in follicles from three 365-day ovaries of different gilts.

### Knockdown of *BMP15* caused premature luteinization and impaired oocyte quality in TGF follicles

We next examined the expression and activation of factors relevant to follicular development. Results firstly confirmed that BMP15 protein abundantly located in both WT oocytes and GCs of primary to pre-ovulatory follicles (Fig. 5A). Both normal and abnormal TGF follicles showed slightly decreased BMP15 protein accumulation in the less degraded GCs than WT. However, markedly reduced BMP15 expression level was noted in deteriorated oocytes of TGF abnormal (TGFA) follicles (Fig. 5A). TGS ovaries exhibited the minimum BMP15 protein level in those primary-like follicles of 110-day TGS ovaries and highly degraded SFs of 365-day TGS ovaries (Fig. 5A). Thus, we speculated that the *in vivo BMP15* interference efficiency was different in transgenic individuals, which was likely responsed for the two TG ovarian phenotypes (TGF and TGS). Besides, TGS ovaries displayed a phenotype of highly degradation, and serious inhibition of follicular development and cellular activity in the arrested SFs, exactly similar to the phenotypes of BMP15 homozygotes mutations sheep(Braw-Tal et al., 1993) and women with POI(Luisi et al., 2015), with which caused female infertility. Hence, we then chiefly focused on the effects of TGF follicles.

**Fig. 5.**
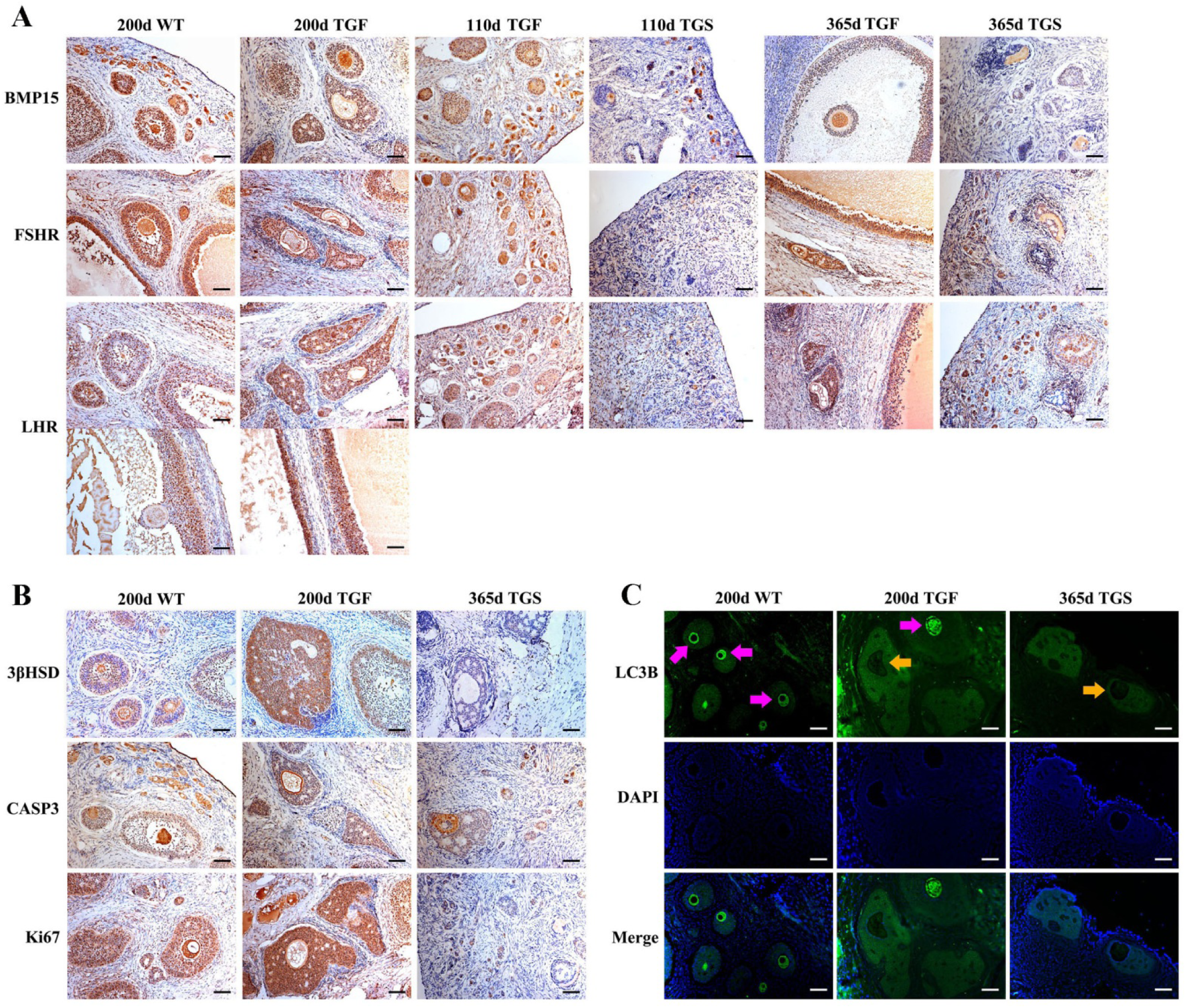
Abnormal TGF follicles showed premature luteinization and impaired oocyte quality. **(A)** Immunostaining on ovarian sections indicates expression of BMP15 decreased remarkably in TGS abnormal follicles, but only slightly reduced in TGF follicles, as compared to WT follicles. FSHR shared a similar expression pattern with BMP15, the expression pattern of LHR was contrary to BMP15. It expressed higher in TGF GCs of both preantral and antral follicles. Scale bar =100μm. **(B)** Follicular cell apoptosis, proliferation, and premature luteinization were evaluated by immunostaining with Caspase3, Ki67, and 3βHSD respectively. Notably higher expression level of 3βHSD was discovered in abnormal TGF follicles. However, expressions of Caspase3 and Ki67 was not significantly different between abnormal TGF follicles and WT follicles. Scale bar =100μm. **(C)** Immunofluorescence images demonstrates intensive expression of autophagy-related protein LC3B in oocytes of normal follicles of TGF and WT ovary, but barely expressed LC3B in oocytes of abnormal follicles of TG (TGF and TGS) ovaries. Purple arrow, oocytes in normal follicles; Orange arrow, oocytes in abnormal follicles. Scale bar= 100μm.

In TGF follicles, we found that the expression patterns of both GDF9 and FSHR, the BMP15 cooperator and down regulator respectively, were corresponding to BMP15 (Fig. S5, Fig. 5A) in TGF follicles. Whenas there were no changes in expression of BMP15 receptors (ALK6 and BMPR2) (Fig. S5). Contrary to BMP15, the luteinizing hormone receptor (LHR) expressed higher in TGF follicles as compared to WT (Fig. 5A). This excess expression of LHR suggested premature luteinization in TGF follicles, which was also demonstrated by the dramatically raised expression of 3βHSD (3β-hydroxysteroid dehydrogenase) in TGF SFs (Fig. 5B)(Grasa et al., 2016). In consideration of the striking features of reduced follicle number and degraded GCs in the TGF follicles, we then detected the expression levels of caspase 3 and Ki67 to assess the cell apoptosis and proliferation activity. Surprisingly, there was no change in expression of both caspase 3 and Ki67 in the degraded TGFA follicles (Fig. 5B). The later investigation of BMP15 mediated signaling pathways suggested an underlying mechanism. We emphasized that notably weakened Smad1/5/8 activity in TGFA follicles when compared to TGF normal SFs (Fig. 6A), but slighter attenuated Smad2/3 phosphorylation was shown in these abnormal follicles (Fig. 6B). It was likely that Smad1/5/8 mainly contributed to the inhibition of follicular development of the TGFA follicles, whereas Smad2/3 activated in a BMP15 independent pathway and played a role in supporting growth of these less degraded follicles. Except for GCs, we also discovered impaired oocyte quality in TGFA follicles. As showed in Fig. 5C, an undetectable level of autophagy-related protein LC3B (microtubule-associated protein 1 light chain 3)(Jiang et al., 2017) was shown in oocytes of TGFA SFs, while oocytes of WT and TGF normal follicles displayed a normal level of LC3B. This result demonstrated that autophagy activity of the oocytes of TGFA follicles was largely weakened, which was fundamental to many oocyte cellular processes(Su et al., 2017).

**Fig. 6.**
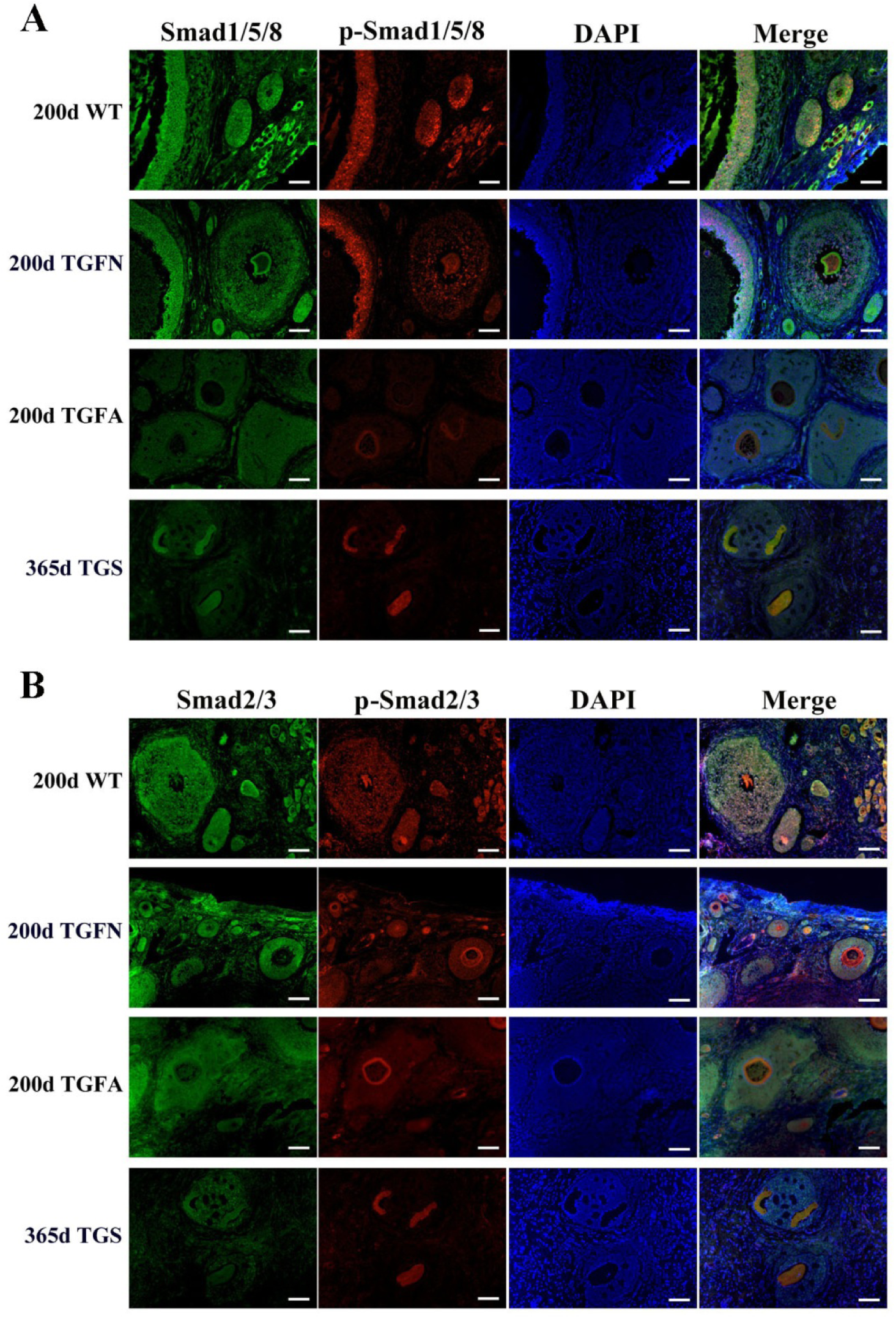
Smad1/5/8 signaling pathway was inhibited in abnormal follicles in TGF ovary. **(A)** Immunofluorescence images showed Smad1/5/8 pathway was evidently less activated in abnormal follicles in TGF ovaries, and severely inhibited in highly degraded 365-day TGS follicles, as compared to that in normal follicle of TGF and WT ovaries. **(B)** Immunofluorescence images demonstrated a mild decrease in Smad2/3 signaling in both normal and abnormal follicles of TG ovaries. Smad2/3 signaling was remarkably inhibited in highly degraded 365-day TGS follicles. TGFN, normal follicles in TGF ovary; TGFA, abnormal follicles in TGF ovary. Scale bar = 100μm.

### Knockdown of *BMP15* resulted in dynamic transcriptomic alteration during TGF follicular growth

To further investigate the regulatory role of BMP15 in porcine follicular development, RNA-seq was carried out on follicles or GCs captured by laser capture microdissection (LCM) method from frozen sections of both WT and TGF ovaries. LCM-captured follicles were categorised to three stage of follicular development: primary follicle (PF), secondary follicle (SF), and small antrum follicle (SAF) stages. For large antrum follicle, only parietal granulosa and thecal cells (APC) were captured by LCM for RNA-seq. The follicles or APCs were captured from frozen sections of each 5 TGF and WT ovaries of gilts at age ranging from 60 to 170 days. The gene profiles of 34,640 genes generated by RNA-seq were used for identification of differentially expressed genes (DEGs), and GO and pathway enrichment analysis basing on intra (between each two continuous follicle stages in either WT or TGF sample) and inter (between each follicle stage of WT and TGF sample) effect comparisons (Table S6). In intra effect comparisons, the largest number of DEGs (3503 DEGs) was found in SAF^WT^/SF^WT^ comparison, and the least number of DEGs (350 DEGs) was found in APC^WT^/SAF^WT^ comparison (Table S6). However, in contrast to WT, during TGF follicular development, the lowest number of DEGs was found in SAF^TGF^/SF^TGF^ comparison, and the largest number of DEGs was found in SF^TGF^/PF^TGF^ comparison (Table S6). Striking difference of the dynamics of transcriptions between WT and TGF follicular development were also found in GO (Fig. 7A) and pathway (Fig. S7A) enrichment. In WT ovary, more DEGs was presented in enriched GO during the dynamical transition from early primary to secondary follicle stage, and from secondary to small antrum follicle stage, but less DEGs was presented in enriched GO during small antrum follicle to large antrum follicle stage transition. Whereas, in TGF ovary, more DEGs was presented in enriched GO during the dynamical transition from small antrum follicle to large antrum follicle stage, but less DEGs was presented during secondary to small antrum follicle stage transition (Fig. 7A). These results in GO enrichment were in line with that found in pathway enrichment (Fig. S7A). Based on the intra effect analysis, it seemed like that the GCs differentiation during the late secondary stage and early antrum formation(Hennet and Combelles, 2012), had been delayed during TGF follicular dynamical development. BMP15 probably played a more important role during the dynamical development of secondary and subsequent follicle stages rather than in early stages.

**Fig. 7.**
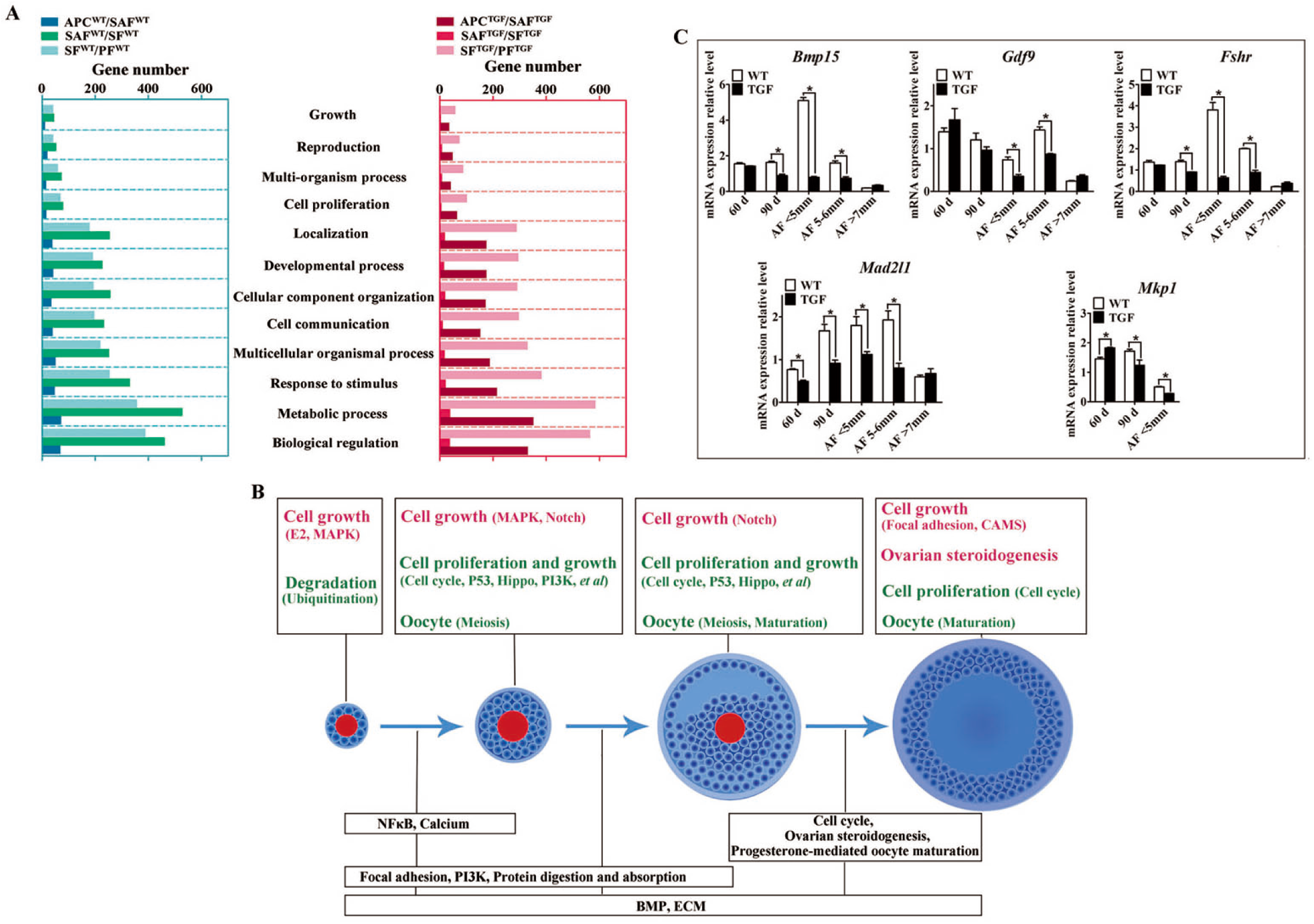
Altered follicle dynamic transcriptomes during TGF follicular development. **(A)** Different number of DEGs was presented in enriched 12 biological processes (GO) between WT and TGF dynamic transcriptomes during follicular development. In TGF ovary, the least DEGs was presented in enriched GO during the transition from secondary to small antrum follicle stage; In WT ovary, the least DEGs was presented in enriched GO during the transition from small antrum to antral follicle transition. The most DEGs was presented in enriched GO during the transition from primary to secondary follicle stage in TGF ovary, whereas in WT ovary, that was presented during the transition from secondary to small antrum follicle stage. Gene Ontology-based analysis was conducted by Webgestalt software. **(B)** Summary of important pathways (based on inter effect analysis) involved in follicular development. Up-regulated pathways are shown in red, and down-regulated pathways are shown in green. **(C)** A subset of DEGs was validated by qPCR. 60 d stands for 60-day ovarian tissue, which was mainly compose of early stage follicles (primordial, primary, and early secondary follicles). 90 d represents for 90-day ovarian tissue, which was mainly composed of secondary follicles without antral follicles. AF<5mm, antral follicle with a diameter <5 mm. AF 5-6 mm, antral follicle with a diameter of 5-6 mm. AF>7mm, antral follicle with diameter >7 mm.

In consideration of the expression and function of BMP15 during follicular development were stage-specific(Paradis et al., 2009; Sun et al., 2010), we next conducted the RNA-seq data analysis based on inter effect comparisons. Both DEGs number and their clustering results revealed distinct effect on knocking down of BMP15 on each follicle stage, where the largest number of DEGs was found in the secondary follicle stage (Fig. S8A). Based on these DEGs, we enriched total 15 up-regulated and 26 down-regulated pathways (Fig. S7B) in all four follicle stages. 10 pathways were enriched during the three dynamical transitions of each two continuous follicle stages, in which DEGs were identified based on the combination of the intra and inter effect comparison (Fig. S8). These pathways were then illustrated in Fig. 7B according to their relevant function. Interestingly, knocking down of BMP15 seemed to lead to an increase of signaling of cell growth in primary follicle, due to the significant up-regulation of Estrogen and MAPK pathways, and the down-regulation of ubiquitin protein degradation pathway. However, knocking down of BMP15 was likely to inhibit GCs proliferation and growth from secondary follicle stage onward, because of the significant down-regulation of pathways including Cell cycle, Hippo, P53, PI3K *et al*, which also implied a potential involvement of BMP15 in these pathways. Though GCs proliferation and growth were inhibited by knocking down of BMP15, the significantly up-regulated MAPK pathways in primary and secondary follicle stages, and Notch pathway in secondary and small antrum follicle stages may play a role for partial compensation on the cell proliferation and growth in preantral follicles to support the continuous development of certain percent of follicles to term. Furthermore, knocking down of BMP15 did not result in significant pathway alteration during the transition from secondary to small antrum follicle stage t (Fig. 7B). Combined with the finding of that the minimum DEGs was presented in SAF^TGF^/SF^TGF^ comparison (Table S6), it suggests that knocking down of BMP15 may cause an inhibition of GCs differentiation and abnormal development in preantral follicles. In large antrum stage, DEGs involved in ovarian steroidogenesis (*Lhr*, *Cyp17, 3βHsd et al*) were significantly up-regulatd in theca cells of TGF follicles (Table S7), possible associated with the undergoing of premature luteinization of TGF follicles (Fig. 5B). Except for GCs, we also enriched significantly down-regulated DEGs involved in oocyte meiosis and maturation in TGF follicles beyond primary follicle stage, possibly related to the impaired oocyte quality (Fig. 5C).

Moreover, we found *in vivo* knocking down of BMP15 resulted in significant decrease of *Bmp15* expression level from primary to small antrum follicle stage (Table S8), which was confirmed by qPCR analysis (Fig. 7C). Through a correlation analysis of the DEGs, we predicted 13 downstream regulated genes of *Bmp15*, in which 6 genes (*Atrx*, *Amd1*, *Dtd2 et al*) were positive correlated, and 7 genes (*Fgf9*, *Igfbp7*, *Cmpk2 et al*) were negatively correlated (Fig. S8B, Table S8). Furthermore, an unexpected significantly decreased expression of *Fshr* in TGF follicles was detected by transcriptomic analysis (Table S7), and confirmed by qPCR (Fig. 7C), which seemed to be inconsistent to the previous perspective that BMP15 played a role in suppression of *Fshr* expression(Abir and Fisch, 2011; McMahon et al., 2008; Otsuka et al., 2001; Shimizu et al., 2019).This discrepancy was probably due to species difference or the abnormal development of TGF GCs, including degradation, premature luteinization *et al*. In addition, mRNA examination revealed that *Mad2l1* (*Mitotic spindle assembly checkpoint protein*) decreased significantly in TGF ovarian tissues and antrum follicles (Fig. 7C), implying the inhibition of TGF cell mitosis. Increased expression of *Mkp1* (*Dual specificity protein phosphatase 1*) in 60-day TGF ovarian tissues but decreased expression of this gene in 60-day TGF ovarian tissues and antrum follicles (Fig. 7C) seemed to be related to the up-regulated MAPK pathway in early TGF preantral follicles (Fig. 7B).

### Knockdown of *BMP15* caused reduced capacity of TGF follicles to ovulate

The evidence of disordered estrous cycle, abnormally enlarged antral follicles, and that no corpus lutein was observed in sexually mature TG gilts until 365 days old, demonstrates that knockdown of *Bmp15* could cause dysovulation. To investigate the underlying factors causing dysovulation by knocking down of BMP15, COCs (oocyte-cumulus complexes) were isolated from antrum follicles with a diameter of 5-7 mm for single-cell RNA sequencing. As expected, sequencing results showed a drastic decreased of *BMP 15* in TGF COCs (Table 1), which was confirmed by qPCR analysis in antrum follicles (Fig. 7C). Interestingly, GDF9, the closely related homologous protein of BMP15, was down-regulated by knocking down of BMP15 (Fig. 7C and Table 1). However, the expressions of another BMP poteins (*Bmp4*, *Bmp6*) were not affected by knocking down of BMP15. Surprisingly, BMP15 receptors (*Bmpr2* and *Alk6*) as well as its signaling protein *Smad8* were significantly up-regulated, which was probably related to the increased expression of *Fst* and *Inhα* or another activin. About four-folds decreased expression of both *Fshr* and *Hsd17β* (Table 1 and Fig. 8C) might contribute to the dramatically reduced E2 production and the absence of dominant follicle selection in TGF antrum follicles. Up-regulated expression of steroidogenesis related factors including *Lhr*, *Star* (*steroidogenic acute regulatory protein*), *Cyp11a* (*cytochrome P450 family 11 subfamily A member*), *Cyp19a* (*cytochrome P450 family 19 subfamily A member*) (Table 1 and Fig. 8B, C) in TGF COCs was consistent with the results of transcriptomic analysis of TGF APC (Table 7), which might contribute to the undergoing of premature luteinization. It has been reported that the down expression of *Amhr2* (Anti-Mullerian hormone receptor type 2) and *Cx43* (Gap junction protein alpha 1) induced by LH in preovulatory follicles was important to ovulation(Norris et al., 2008; Pierre et al., 2013). Thus the increased expression of these two genes in TGF COCs potentially resulted in a decreased capacity of oocyte meiosis resumption and ovulation. In addition, markedly decreased expression of oocyte quality related genes (*Bmp15*, *Gdf9*, *Zp2*, *Zp3*, *Zar1, and Irf6*) strongly implied a reduced oocyte competence in TGF COCs.

**Table 1.**
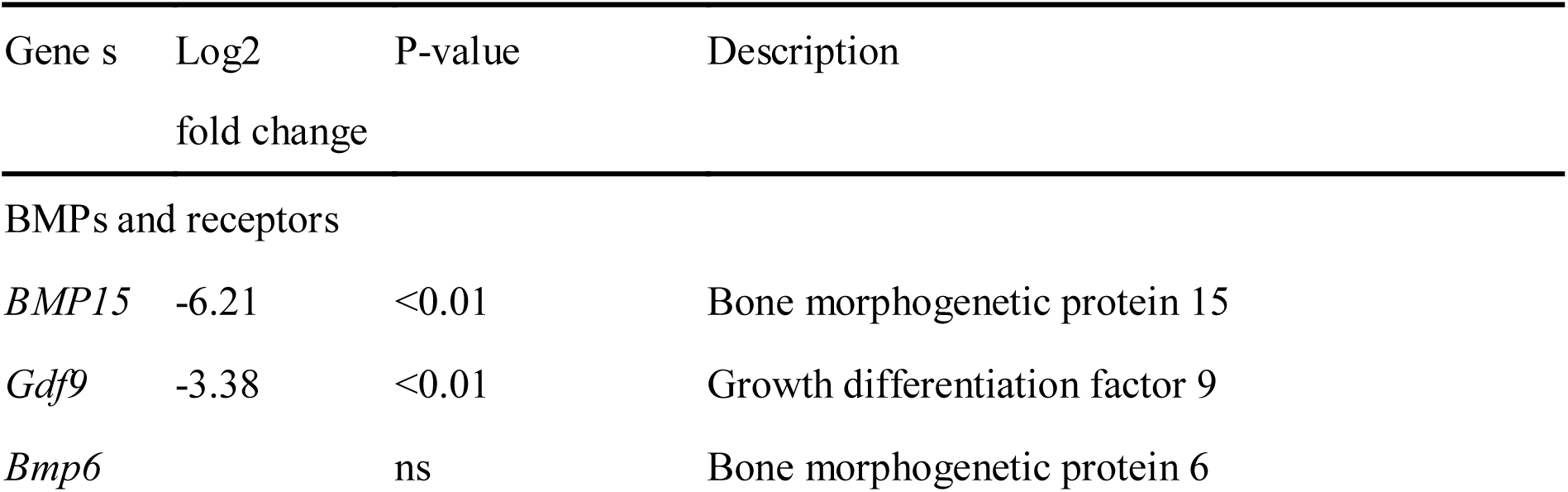
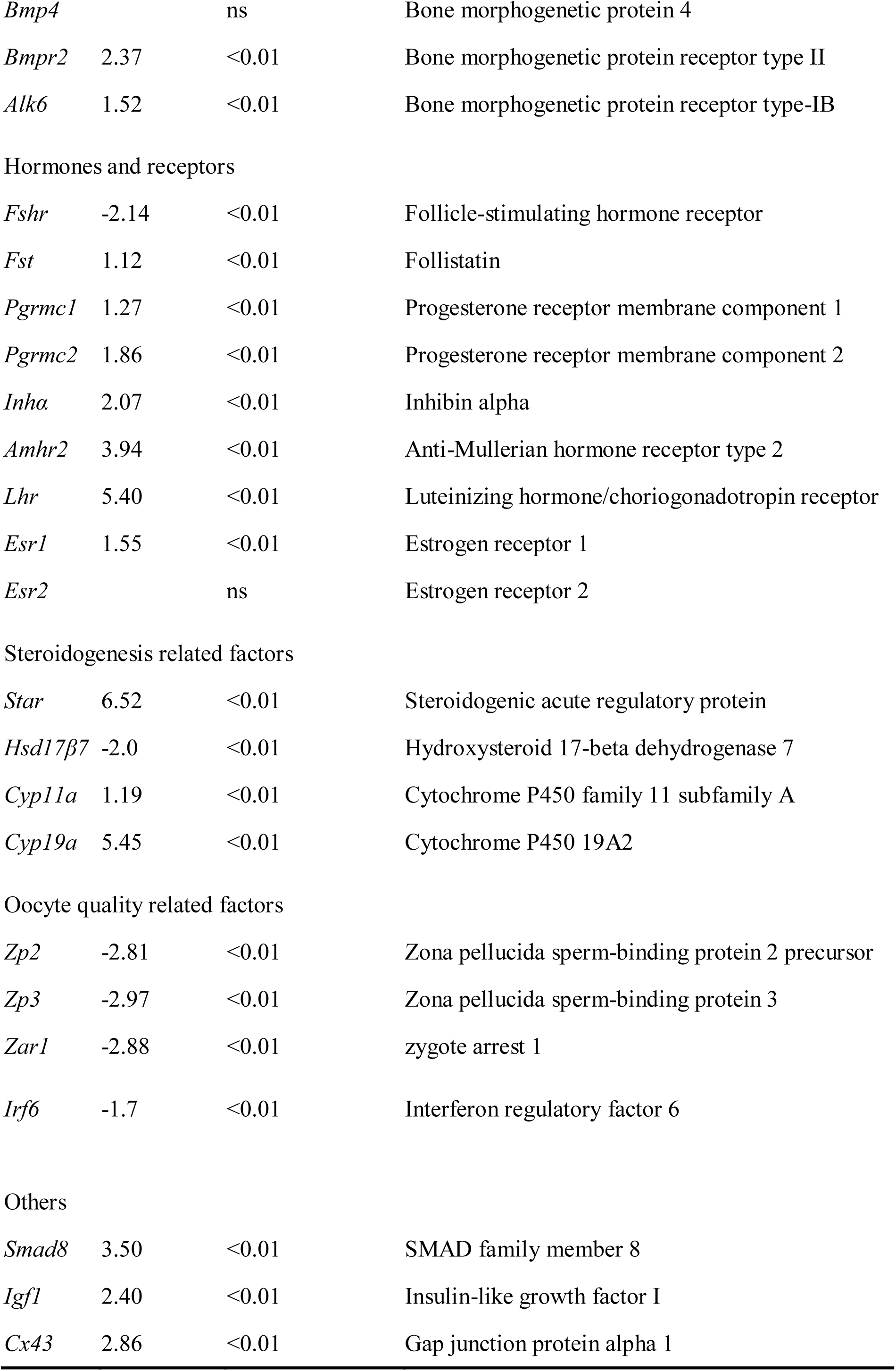
Interest genes that related to COCs function.

**Fig. 8.**
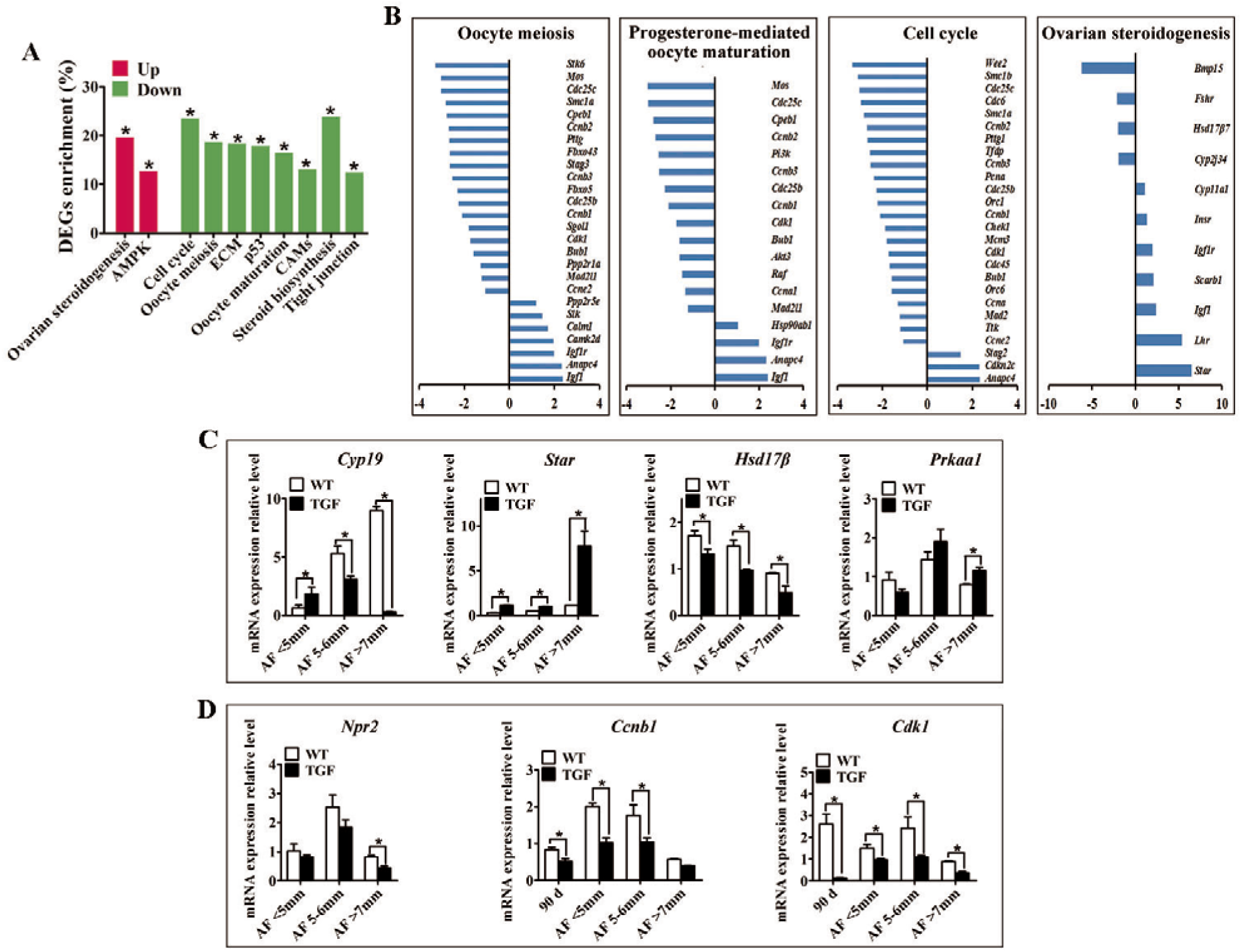
Single cell RNA-seq on TGF COCs showed affected process in ovulation. **(A)** Pathway enrichment revealed two significantly up-regulated pathways which potentially contributed to increased number of large antral follicles, and 8 significantly down-regulated pathways that are involved in cumulus cell function and oocyte maturation. **(B)** Pathways of Oocyte meiosis, Progesterone-mediated oocyte maturation, Ovarian steroidogenesis, and Cell cycle, which are closely related to oocyte maturation and ovulation, showed DEGs enrichment of 22.4%, 21.2%, 25.5%, 22.6% respectively. **(C)** qPCR validation of mRNA expression level of DEGs involved in the pathways of Ovarian steroidogenesis (*Cyp19*, *Star*, *Hsd17β*) and AMPK (*Prkaa1*). **(D)** Quantification of mRNA level of genes related to oocyte meiosis indicated decreased expression of MPF, but unaffected expression of *Npr2* during TGF follicular development. 90 d represents for 90-day ovarian tissues.

Total 2,820 DEGs (885 up-regulated, 1,935 down-regulated) was generated for pathway enrichment. The significantly up-regulated AMPK (Fig. 8A, C) and Ovarian steroidogenesis (Fig. 8A, B) pathways were likely to contribute to the greater number of large antrum follicles in TGF ovaries, according to the findings of previous studies in sheep(Foroughinia et al., 2017) and sow(Knox, 2005). However, pathways (Cell cycle, P53 *et al*) involved in cell proliferation and growth, were significantly down-regulated (Fig. 8A, B), which was consistent with the results of the dynamic transcriptomic analysis of TGF follicles (Fig. S7B). Furthermore, four pathways including oocyte meiosis, oocyte maturation, cell cycle, and ovarian steroidogenesis, which were closely involved in regulation of oocyte maturation and ovulation, presented a DEGs enrichment more than 20% (Fig. 8A,B). In total, these results revealed both impaired function of cumulus cells and oocyte maturation in TGF COCs, suggesting a reduced capacity to ovulate.

However, surprisingly, the expression of *Impdh* (*Inosine monophosphate dehydrogenase 2*) and *Npr2* (*Natriuretic peptide receptor 2*) was not affected (Fig. 8D and Table S9). These two genes have been reported in mice to be up-regulated by BMP15 and GDF9 during the activation of maturation promoting factor (MPF) (Cyclin B and CDK1) and stimulation of oocyte meiotic resumption *in vitro*(Wigglesworth et al., 2013). Instead, we discovered significantly decreased expression of *Cyclin B* and *Cdk1* (Cyclin dependent kinase 1) in TGF follicles from secondary stage onward (Fig. 8D and Table S9), which implies the involvement of BMP15 in modulating porcine oocyte meiosis possibly through regulating the expression of MPF.

## Discussion

The effect of BMP15 mutations on altering ovarian follicular development and ovulation rate was firstly discovered in Inverdale (FecX) sheep(Braw-Tal et al., 1993; Davis et al., 1992; Smith et al., 1997). In these sheep, ewes with single allele of inactive *BMP15*gene showed increased ovulation rate and a higher incidence of twin or triplet births, while ewes with bi-alleles of inactive *BMP15* gene were sterile with primary ovarian failure phenotype(Galloway et al., 2000). Studies on animals immunized with different regions of BMP15 peptide revealed that an increased ovulation rate can be found in females with BMP15 being partially neutralized, but an inhibition of follicular growth and ovulation was found in females with vast majority of active BMP15 being neutralized(McNatty et al., 2007). Therefore, different extent of reduction of biologically active BMP15 protein level seems to lead to varied effect on fertility. In this study, we found two different ovarian phenotypes (TGS and TGF) in our *BMP15* knockdown gilts. The different ovarian phenotypes might be caused by different *in vivo* expression level of BMP15.Indeed, in TGS ovaries, the marked reduced level of both BMP15 mRNA (Fig. 1E) and protein (Fig. 5A) revealed that the majority of BMP15 was knockdown by integrated shRNA. In contrast, in TGF ovaries, the BMP15 protein accumulated abundantly, and was only slightly lower than that in WT ovaries (Fig. 5A), despite the mRNA level of BMP15 had decreased to the half of wild-type as detected in 365-day (Fig. 1E) and 30-day (Fig. 1F, G) ovarian tissues. Therefore, the less interference of BMP15 expression in TGF ovaries may confer them a less severely impaired ovarian phenotype, as TGF ovaries contained each stage of follicle, but presented remarkably reduced follicle number, increased ratio of abnormal follicle, disordered reproductive hormones, and ovulation dysfunction. We found that TGF and TGS ovaries could concurrently appear in single TG gilts (Fig. 3A). The difference in *in vivo* interference efficiency of integrated shRNA plasmid between bilateral ovaries in single TG individual may be caused by unknown complicated regulatory mechanism of transgene expression, possibly including epigenetic factors.

Previous reports in sheep revealed that heterozygous mutations in BMP15 had been proved to inhibit GCs growth but increase GCs sensitivity to FSH, leading to increased ovulation of smaller matured follicles with reduced amounts of E2 and inhibin (Fabre et al., 2006a; Otsuka et al., 2001; Otsuka et al., 2000; Shackell et al., 1993). However, this was inconsistent to our results. As the *in vivo* mRNA level of BMP15 in TGF ovaries was knocked down to half of wild-type in TG gilts, thus these TG gilts with TGF ovaries could be considered as pigs with heterozygous mutations in BMP15. In different to ewes heterozygous for mutations in BMP15, our TG gilts did not present increased ovulation rate, but in contrast, a dysfunction in ovulation, as corpus lutein can not be found in TGF ovaries from TG gilts of age younger than 365 days, though they can be observed in TGF ovaries from TGF gilts at age of 400 and 500 days (Fig. S4A), indicating a delayed ovulation in TGF gilts. In addition, significantly decreased amount of total and smaller antral follicles in TGF ovaries (Fig. 4A, B) seemed incapable to support an increased ovulation rate. Moreover, the appearance of abnormally enlarged antral follicles with lower FSHR expression level and E2 production (Fig. 4) may indicate that TGF follicles couldn’t ovulate at normal size probably due to the attenuated FSH sensitivity and a lack of dominant follicle selection in TG gilts. Besides, premature luteinization of GCs possibly caused by insufficient FSH stimulation also may contribute to the dysovulation of TGF gilts. Thus, these results in TGF puberty gilts were also different from the poly-ovulatory *Bmp15^−/−^* mice, which showing normal follicular development and could ovulat at puberty(Yan et al., 2001). Hence, we may suggest the importance of BMP15 in regulating ovulation was species-specific different, that not only between mono- and poly-ovulatory mammals, but also between poly-ovulatory species.

BMP15 has been proved to suppress *Fshr* expression in GCs to affect GCs proliferation and steroidogenesis in antral follicles of rodent(McMahon et al., 2008; Otsuka et al., 2001) and human(Abir and Fisch, 2011; Shimizu et al., 2019). Previous studies on sheep indicated that heterozygous mutations in BMP15 could increase the sensitivity of GCs in antral follicle to FSH stimulation, leading to increased ovulation rate(Fabre et al., 2006a). However, a recent study showed that treatment with BMP15 caused increased expression of *Fshr* in bovine preantral follicles after 12 days culturing(Passos et al., 2013). These conflicting reports probably caused by species-specific differences and the different response to BMP15 stimulation in each follicular development stages. In this study, we found that, in different to the results found in sheep, knockdown of *BMP15* did not increase but significantly inhibited *Fshr* expression in both preantral and antral follicles (Fig. 7E, Table 1, Table S7). Furthermore, the findings of inhibition in GCs proliferation and differentiation, increased expression of genes involved in steroidogenesis (*StAR*, *Cyp11a*, 3*β*HSD) (Fig. 5B, 8C, Table 1, Table S7), drastically decreased of E2 production (Fig. 4D), subsequent absence of dominant follicle selection in the TGF follicles, were likely to be consequences of the declined sensitivity of GCs to FSH. Suppression of *FSHR* expression in preantral follicles of TG gilts implies that BMP15 could stimulate porcine follicle growth and development in an earlier follicle stage rather than gonadotropin dependent period(Mori, 2016). Given the degradation of GCs and abnormal structure GCs layers observed in TGF ovaries, another possible reason for the decreased expression of FSHR might be related to the impaired GCs development caused by BMP15 deficiency.

As a paracrine and autocrine factor of oocyte, BMP15 can promote not only the development of follicular somatic cells, but also the development of oocyte itself. Studies on the oocyte *in vitro* maturation have demonstrated BMP15 were capable to stimulate cumulus cell expansion(Braw-Tal et al., 1993; Lin et al., 2014; Peng et al., 2013; Sudiman et al., 2014; Sugiura et al., 2010), promote signalings of LH-induced maturation of the cumulus-oocyte complex(Su et al., 2010), improve oocyte quality(Caixeta et al., 2013; Hussein et al., 2006), and increase blastocyst rate and embryonic development of fertilized oocytes(Gode et al., 2011; Wu et al., 2007). Recently, the expression level of BMP15 has been suggested as a diagnostic marker of oocyte quality(Wu et al., 2007). In this study, we confirmed the important role of BMP15 in porcine oocyte development *in vivo*. We found that knockdown of BMP15 could cause oocyte degeneration or abnormal enlargement (Fig. 3G, H), and lacked of normal autophagy activity in oocytes of abnormal preantral follicles (Fig. 5G). Further transcriptomic analysis of the follicle and COCs also implies genes and pathways involved in oocyte meiosis and maturation were affected in TGF follicles from secondary stage onward (Fig. 7B, 8A). Previous studies have indicated possible underlying mechanisms of BMP15 in regulating oocyte meiosis. One study considered that BMP15 and GDF9 can promote oocyte meiotic resumption in mice through up-regulation of *Npr2* and *Impdh*(Wigglesworth et al., 2013). Another study showed that inhibiting BMP15 signaling pathway by Smad2/3 phosphorylation inhibitor resulted in significantly decreased expression of *Cdc2* and *Cyclinb1* during porcine oocyte *in vitro* maturation(Lin et al., 2014). However, our transcriptomic results showed that the expression level of *Npr2* and *Impdh* both were not affected in both TGF follicles and COCs, instead, expression level of MPF (*Cdc2* and *Cyclinb1*) decreased significantly (Fig. 8D and Table S9). Therefore, our results might support an und**e**rlying mechanism of BMP15 involved in porcine oocyte meiosis and maturation through regulating the expression of MPF, however, this requires further studies to elucidate.

In summary, knockdown of *BMP15* caused markedly reduced fertility of TG gilts mainly through inhibition of both GCs and oocyte development (Fig. 9). The suppression of GCs proliferation and differentiation led to decline in number of early follicles, GCs degradation, and reduced sensitivity of GCs to FSH stimulation with consequence of premature luteinization, higher LHR expression, but lower E2 production in large antral follicles. The effect on oocyte development directly led to impaired oocyte quality and oocyte meiotic maturation. Consequently, large antral follicles abnormally enlarged resulting in dysovulation and disordered reproductive cycle hormones. Our results revealed a remarkable physiological suppression of porcine ovarian follicular development and ovulation in *BMP15* knockdown gilts, demonstrating an essential role of BMP15 on porcine female reproduction, and providing new insights into the regulatory role of BMP15 in poly-ovulatory mammals. Our findings provided important implications on further investigation of the complicated regulatory function of BMP15 in female fertility of poly-ovulatory species, and development of possible strategies for improving porcine female fertility through modulation of BMP15 expression.

**Fig. 9.**
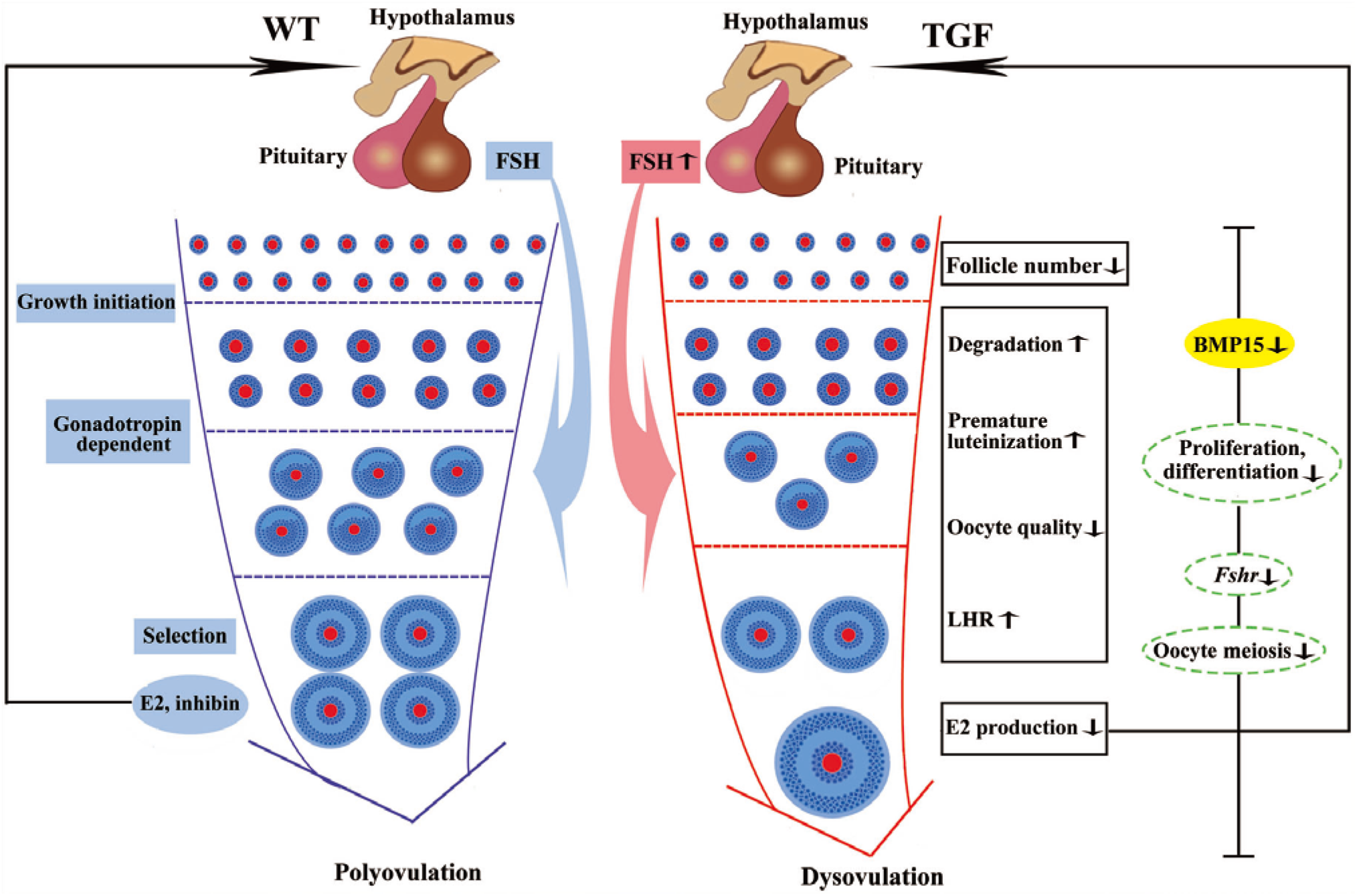
Summary of biological functions and possible regulatory mechanism of BMP15 in TGF follicular development. As compared to WT gilt, knocking down of BMP15 caused remarkable follicle number reduction during the TGF follicular developmental process, accompanied by impaired oocyte quality, degradation, and premature luteinization in TGF preantral follicles, which may leads to increased expression of LHR and dramatically decreased E2 production in TGF antral follicles, resulting in lack of dominant follicle selection but abnormally enlarged antral follicles, presenting an dysovulation phenotype. Decreased E2 production in TGF antral follicles may have a negative feedback to pituitary to increase the expression of *Fsh*. However, the decreased expression of *Fsh* receptor *Fshr* in GCs beyond primary follicle stage by knocking down of BMP15 may attenuate its function on stimulation the growth of antral follicles. In addition, knocking down of BMP15 leads to inhibition in cell proliferation and differentiation, declined of oocyte quality and meiotic maturation during the TGF follicular development, these together contributes to the appearance of enlarged follicle in TGF ovary and dysovulation. Colored ellipses represent results based on morphological observation and molecular examination. Dotted ellipses represente results based on the transcriptomic analysis.

## Methods

### Construction of shRNA expression vectors and evaluation of shRNA interference efficiency

Five shRNAs (Table S1) targeting to porcine *Bmp15* mRNA was designed and selected by Invitrogen’s web-based siRNA design software (https://rnaidesigner.invitrogen.com/rnaiexpress/). Human U6 promoter followed by each shRNA sequence was individually synthesized (Sangon Biotech, China), and cloned downstream of the EGFP expression cassette on pEGFP-N1 vector (Takara Bio, USA) to generate each pEGFP-*Bmp15-*shRNA expression vector (Fig. 1A). Meanwhile, a scramble shRNA expression vector generated as negative control. To evaluate the RNA interference efficiency of shRNA, porcine *Bmp15* CDS was synthesized (Sangon Biotech, China), and cloned into psiCheckIIvector (Promega, USA) to generate psiCheckII-*Bmp15* plasmid. Each pEGFP-*Bmp15-*shRNA plasmid then was respectively co-transfected with psiCheckII-*Bmp15* plasmid into HEK293 cells. After 48 h culturing, transfected cells were collected, and subjected to RNA interference efficiency detection by using a dual-luciferase reporter system (Promega, USA). The shRNA with most efficient RNA interference efficiency then was selected for generation of BMP15 knockdown pig model.

### Generation of *Bmp15* knockdown pig model

Procedures for generation of the *Bmp15* knockdown gilts were illustrated in Fig. S1. Briefly, the selected pEGFP-*Bmp15* shRNA plasmid was transfected into PEFs derived from a male Yorkshire pig. After G418 selection and fluorescence examination, EGFP positive PEFs were used as donor cells for somatic cell nuclear transfer (SCNT). For SCNT, oocytes were recovered by aspirating ovaries collected from abattoir with a 20 G needle connected to syringe, and then cultured in HEPES-buffered tissue culture medium 199 and later maturation medium, until *in vitro* maturation. SCNT by handmade cloning and embryo transplantation were carried out by BGI Ark Biotechnology company, China. After 114 days of pregnancy, we obtained two surviving F0 generation *Bmp15* knockdown transgenic (TG) males. Then we mated one TG boar with wild-type sows through artificial insemination (AI), and obtained F1 generation TG gilt for this study. Sibling gilts without pEGFP-*Bmp15*-shRNA integration were used as controls (WT) in this study. The protocol of animal study was approved by the Institutional Animal Care and Use Committee (IACUC), Sun Yat-sen University (Approval Number: IACUC-DD-16-0901).

### Tissue collection

A total of 54 animals including 25 WT and 29 TG gilts were sacrificed at ages of 30 to 500 days. Among these TG gilts, 6 gilts contained 8 TGS ovaries at age of 110, 160, 200 and 365 days. Tissues of ovary, pituitary, muscle, liver, kidney, heart and uterus were collected. Tissues used for RNA extraction were directly soaked in Trizol reagent (Promega) and frozen in liquid nitrogen quickly. Muscle tissues used for DNA extraction together with the 30-day ovarian tissues used for protein detection, were directly frozen in liquid nitrogen before transported to the laboratory. All the other ovaries were washed in sterilized saline water and photographed. Some of them were weighed later. Each 6 ovaries at age of 60 and 90 days were used for ovarian tissues mRNA detection. And each six 365-day ovaries of WT and TGF gilts were used for ovarian tissues and isolated follicle mRNA detection respectively. Primers for qPCR analysis were shown in Table S3. 10 ovaries (5 WT and 5 TGF) at age of 60 to 170 days were frozen in OCT and stored at −80°C before laser capture microdissection (LCM). Six ovaries (3 WT and 3 TGF) from different 365-day gilts were used for dissection and follicular fluid collection. Each 3 WT and TGF 365-day ovaries were used for COCs collection and later single-cell sequencing. The rest ovaries (24 WT, 24 TGF and 8 TGS) at age of 30 to 400 days were used for HE observation and IHC analysis, they were fixed in 10% (w/v) paraformaldehyde / 0.02 M PBS (pH 7.2) on ice before transportation to the laboratory.

### Identification and characterization of transgenic gilts

Tansgenic pigs were first screened by GFP fluorescence on toes under sunlight, and confirmed by PCR analysis (Table S2) of the integration of pEGFP-*Bmp15* shRNA plasmid in genome using genomic DNA extracted from muscle tissues. The copy number of integrated plasmid was determined by qPCR and Southern blot analysis. mRNA expression level of *Bmp15* in transgenic pigs was detected by qPCR in 365-day ovaries, and BMP15 protein level was detected by Western blot analysis of 30-day ovarian tissues.

### F1 gilts estrous checking and hormone assays

About 50 F1 TG gilts at age of 170 to 400 days were checked daily for signs of oestrus in the presence of an intact mature boar. Each two TG and WT gilts at age about 365 days were chosen for daily vaginal smears analysis, and daily jugular venous blood collection at 9:00 to 11:00 AM for 24 days continuously. Vaginal cell smears analysis and estrous identification were performed as described in a previous reporr(Mayor et al., 2007). Daily blood samples were centrifuged at 1500 g for 15 min, then the serum samples were collected and stored at −80°C. These serum samples were thawed on the ice in prior to be used for quantification of the concentration of oestradiol (E2) and progesterone (P4) by chemiluminescence immunoassay (CLIA) (Siemens, Germany).

### Histological examination

Ovaries derived from gilts at age of 30 to 400 days were fixed in 10% (w/v) paraformaldehyde with 0.02 MPBS (pH 7.2) at 4 °C for about 2 h. Then were cut vertical slices in about 0.5 cm thickness and fixed in fresh 10% (w/v) paraformaldehyde until total 24 h. These slices were mounted in paraffin, and serially cut into 5 μm-thick sections at last by Rotary Microtome (MICROM, Germany), and stained with hematoxylin and eosin (HE). Ovarian HE sections were observed and photographed under a fluorescent microscope (Zeiss, Germany).

Immunohistochemistry (IHC) detection was performed by using the anti-Rabbit HRP-DAB Cell and tissue staining kit (R&D, CTS005) and anti-Goat HRP-DAB Cell and tissue staining kit (R&D, CTS008). Immunohistofluorescence examination was performed by using TSA plus Fluorescein (Perkinelemer, NEL741001KT) and Cyanine3.5 (Perkinelemer, NEL763001KT) kit. Antibodies of BMP15 (Eterlife, EL166380), GDF9 (Eterlife, EL910881), FSHR (Eterlife, EL912710), LHR (Eterlife, EL904141), Caspase3 (Abcam, ab13847), 3βHSD (Abcam, ab154385), p-Smad1/5 (CST, 9516), Ki67(Abcam, ab15580) and LC3B (Arigo, ARG55799) were diluted 1:100 with PBS. While -other antibodies including ALK6 (Santa cruz, sc5679), BMPR2(Santa cruz, sc5683), Smad2/3 (Santa cruz, sc8332), Smad1/5/8 (Santa cruz, sc6031R), and p-Smad2/3 (Santa cruz, sc11769) were diluted 1:50 in PBS.

### Ovary dissection and follicular fluid collection

Each three 365-day TGF and WT ovaries derived from different individuals, were flushed with sterilized saline water and placed in the incubator at 38°C during transportation to the lab. Later, visible antral follicles in these ovaries were dissected by scalpel blade and tweezers, and classified into 3 groups (1–3 mm, 3–5 mm, >5mm) according to their diameter, which was measured by a vernier caliper. Total follicle number of each group was counted. Then, follicular fluid from antral follicles with diameter of 3-5mm and diameter >5mm were collected by a dispensable 10 mL syringe. The concentrations of FSH, LH, E2, and P4 in follicular fluid were quantified by the CLIA method (Siemens, Germany).

### Laser capture microdissection (LCM)

A total of 10 ovaries from each five WT and TGF gilts at age of 60 to 170 days, were embedded in OCT and placed on a cryostat (MICROM, HM560, Germany). All ovaries were cut into 7 μm-thick sections and mounted on RNAse free membrane slides (MMI, 50102). These membrane slides then were fixed in ice-cold 95% ethanol for 1 min, and later washed in 75% ethanol for 30 sec. Afterward, sections were stained following the methods as previously reported(Golubeva et al., 2013). Briefly, staining mixture was prepared with 1% cresyl violet in absolute ethyl alcohol, EosinY, RNAse free water, and 100% ethanol at the ratio of 3:1:4:4. Membrane slides were stained in this fresh staining mixture for 30 sec, then dehydrated through 100% ethanol 1 min three times, and followed 30 s incubation in xylene. Slides were finally dried for 5 min by a hair dryers blowing cold wind, and stored at −80°C until used.

The follicles were distinguished from each other as follows: primary follicle (PF) was defined by a clear monolayer of cuboidal granulosa cells; secondary follicle (SF) was defined by more than two layers of granulosa cells but without any antrum; small antrum follicle (SAF) was defined by obvious small antrum but not completely separated granulose and cumulus cells; antral follicle (AF) was characterized by a big single central antrum and completely separated granulosa and cumulus cell layers. Entire PF, SF, and SAF, but only parietal granulosa and theca cells of AF (named APC) were isolated by LCM. Each types of follicle on 10 sections of each ovary were dissected under 20× magnification microscopic visualization using MMI Cell Cut Plus system (MMI, Swiss). Later, the dissections were treated with 100 μL of TRK Lysis buffer of the MicroElute total RNA kit (Omega) and 2 μL 2-mercaptoethanol. Both TGF and WT lysates were respectively mixed according to their follicle stage after 10 min lysis at room temperature, and stored on dry ice until RNA extraction. A total of 8 LCM-derived RNA samples, including PF^WT^, SF^WT^, SAF^WT^, APC^WT^, PF^TGF^, SF^TGF^, SAF^TGF^, APC^TGF^, were used for transcriptomic analysis. RNA-seq was performed on an Illumina HiSeq2000 using Illumina TruSeq SBS kit v2 (209 cycles including index) to obtain paired-end reads (2×100 bp).

### Single-cell RNA sequencing on COCs

COCs were aspirated from large antral follicles (diameter about 5-7 mm) of each three 365-day TGF and WT ovaries derived from different gilts by using a 20-gauge needle fixed to a 10 mL disposable syringe. COCs then were pooled respectively, and placed on a stereomicroscope (Nikon). Those COCs with several layers of cumulus cells and uniform cytoplasm were selected. Each 10 selected COCs from TGF and WT ovaries was used for RNA micro-extraction by MicroElute total RNA kit (Omega). Total RNA was pre-amplified by SMARTer® Ultra™ Low RNA Kit (Clontech), and sequenced on Illumina Hiseq 2000 sequencing system.

### Analysis of RNA-Seq data

Raw RNA-Seq clean reads were obtained by removing reads containing low quality reads and/or adaptor sequences from raw reads and mapped to the pig genome (Sus scrofa 10.2), allowing up to two base mismatches. Differential expression analysis was performed using the Benjamini approach, genes with an adjusted P value<0.05 and <log2 expressed (DEGs). DEGs lists were submitted to the databases of Novogene company (China) for further enrichment analysis. GO analysis was performed by Webgestalt software. In all tests, P values were calculated using the Benjamini-corrected modified Fisher’s exact test, and P<0.05 was taken as a threshold of significance. Venn diagrams were drawn using the web tool (http://bioinformatics.psb.ugent.be/webtools/Venn/). Gene co-expression analysis declared correlation coefficient at 0.98 as the threshold value. Closely correlated genes then were imported in cytoscape software to generate the co-expression network.

## Acknowledgements

This work was jointly supported by National Transgenic Major Program (2016ZX08006003-006), National Key R&D Program of China (2018YFD0501200), Natural Science Foundation of Guangdong Province (201 6A030313310).

## Ethics approval

All procedures were performed in strict accordance with the recommendations of the Guide for the Care and Use of Laboratory Animals of the National Institutes of Health. The protocol was approved by the Institutional Animal Care and Use Committee (IACUC), Sun Yat-sen University (Approval Number: IACUC-DD-16-0901).

---

**Fig. S1.**
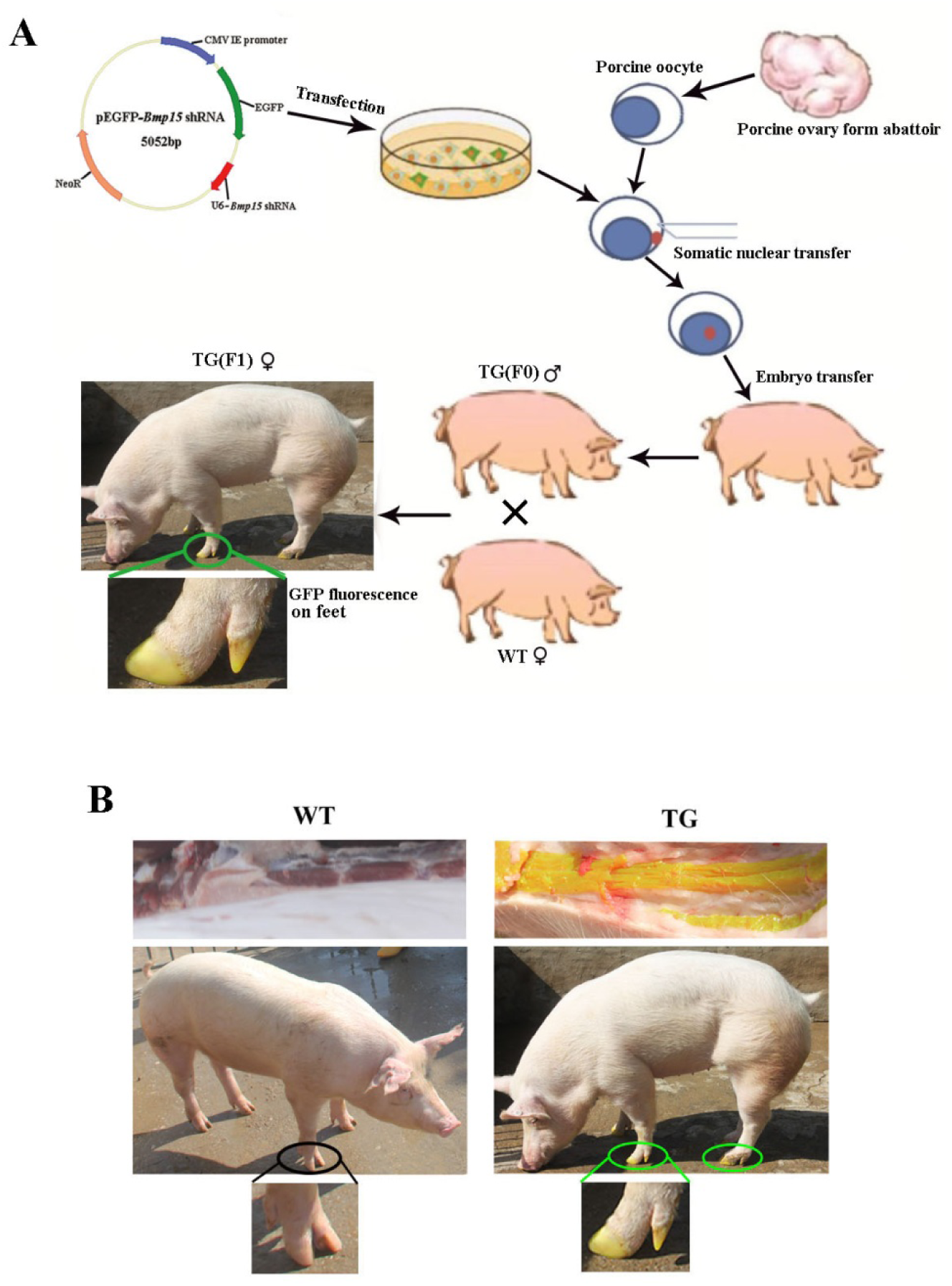
Schematic diagram of generation ofthe *Bmp15* knockdown pig, and identification of transgenic pigs through EGFP fluorescent signal. **(A)** Firstly, we transfected the constructed pEGFP-*Bmp15* shRNA plasmid into Yorkshire PEFs. Then these transfected PEFs were screened with G418 cultured and fluorescence selection to prepare donor cells for somatic cell nuclear transfer (SCNT). Later, we recovered the recipient porcine oocytes by aspirated ovaries from the abattoir. SCNT and subsequent embryo transfer into Large White sow were followed the operation procedure of BGI Ark Biotechnology, China. We at last obtained two healthy neonatal F0 generation transgenic males. One TG boar was mated with wild-type sows to generate F1 gilts. Both F0 and F1 TG pigs showed visible intense GFP fluorescence on toes while subjected to sunlight. **(B)** TG gilts showed remarkable visible GFP fluorescence in muscle and toes under sunlight.

**Fig. S2.**
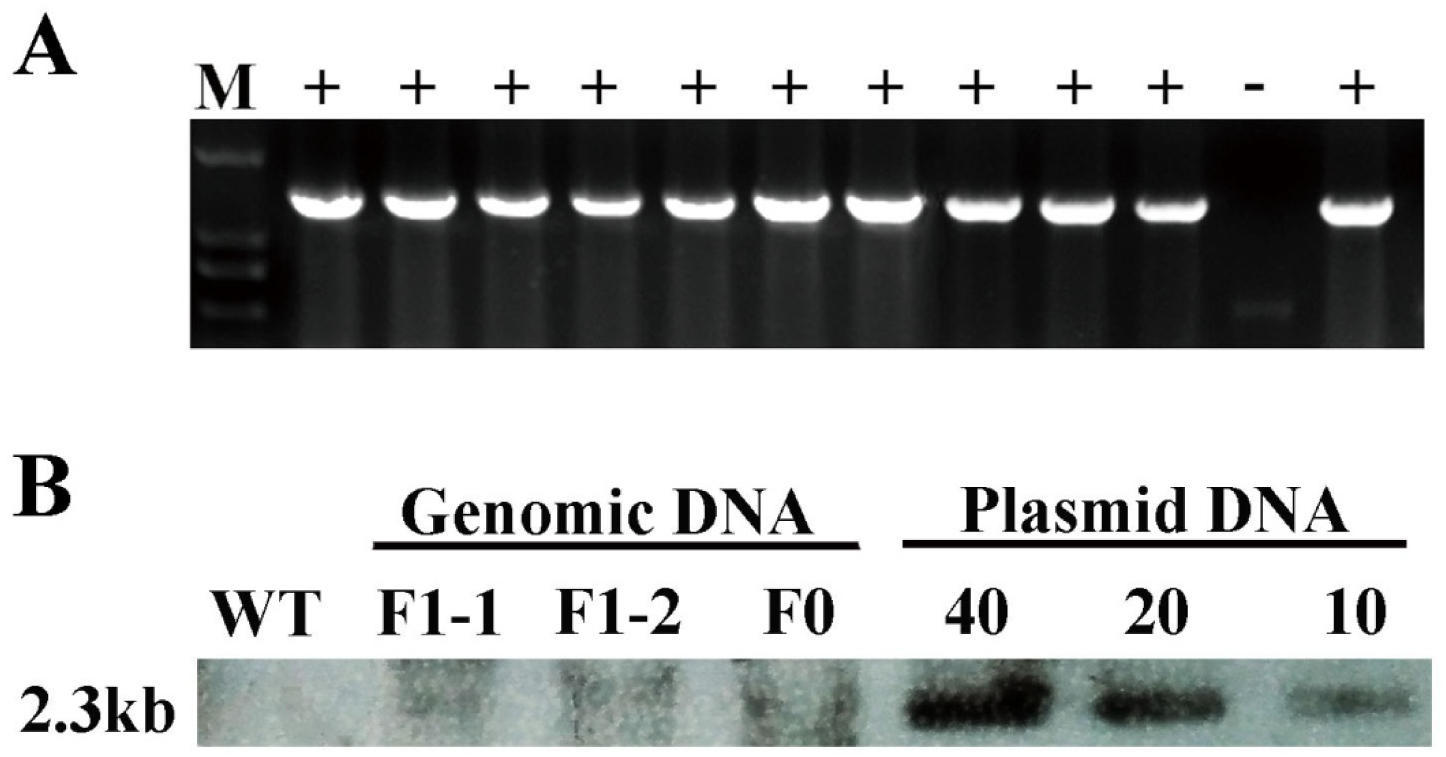
pEGFP-*Bmp15* shRNA plasmid was integrated in genome of TG F0 boar and inherited to TG F1 gilts. **(A)** PCR analysis of the muscle tissue proved that pEGFP-*Bmp15* shRNA plasmid had been transmitted to F1 gilts. +, TG gilt; -, WT gilt; M, DNA Maker. **(B)** Southern blot analysis showed slightly less than 10 copies of constructed plasmids integrated in both F0 and F1 TG pigs, which was consistent with the result of about 7 copies of qPCR analysis (data not shown). DNA with pEGFP-*Bmp15* shRNA plasmid copies of 10, 20, and 40 were used as the positive control.

**Fig. S3.**
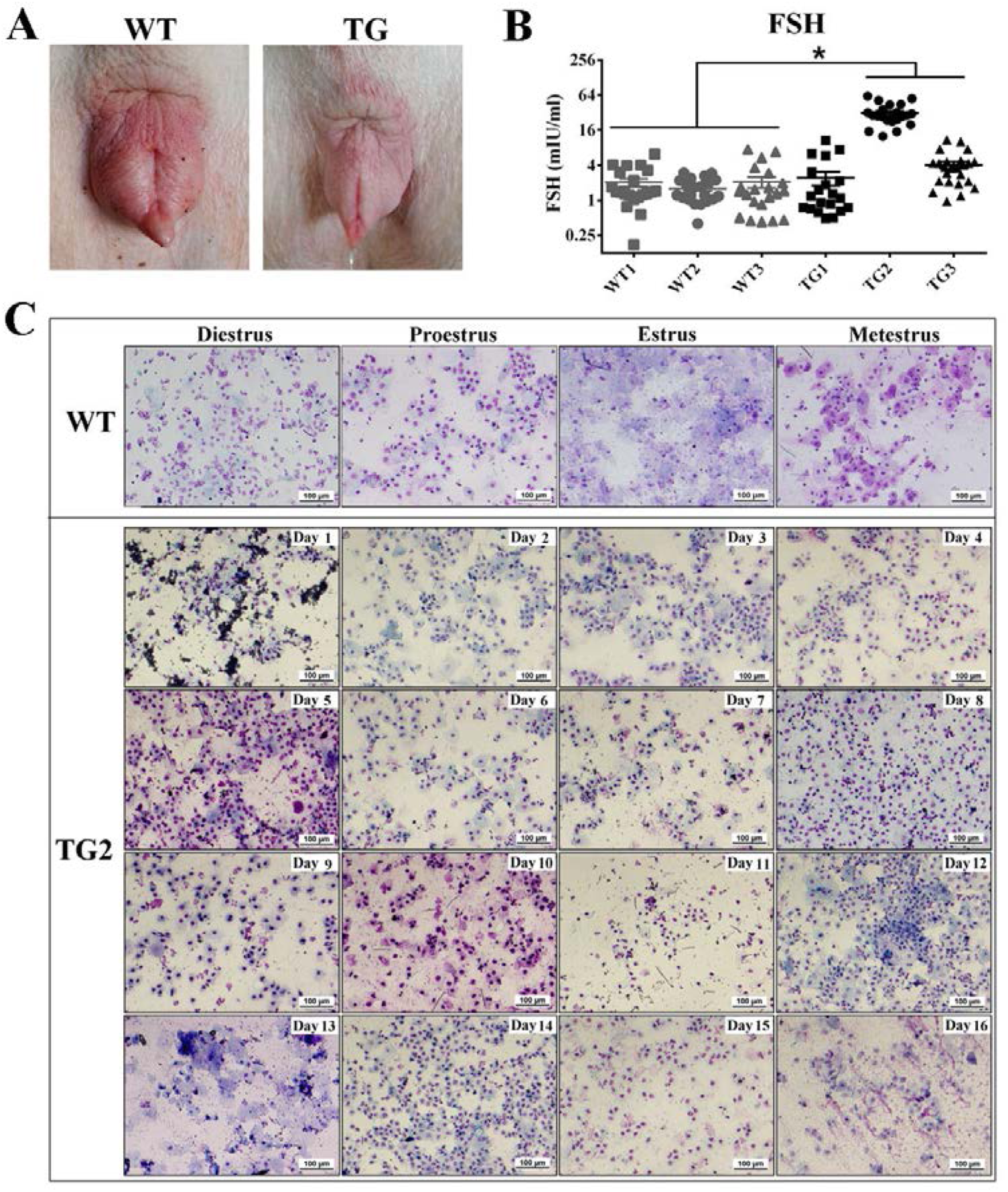
TG gilts didn’t show obvious vulvar appearance change and typical cytologic changes through the estrous cycle, but presented higher FSH concentration in serum. **(A)** Vulvar appearance change (increased redness and swelling) was observed in WT gilt at estrous period, but not observed in 365-day old TG gilts, though they were daily induced by an intact mature boar since 170 days old. **(B)** Two of the three 365-day TG gilts showed higher serum FSH concentration. Serum FSH concentration was measured at a 24h interval for 24 days**. c** Estrous cycle was evaluated by vaginal smears cytology analysis of the 365-day gilts. It was divided into 4 distinct stages according to the appearance and the relative proportions of leucocytes, basal, parabasal, and superficial cells. In WT gilts, proestrus, estrus, metestrus, and diestrus stage were clearly determined through the Giemsa-stained vaginal smears. However, TG gilts presented disordered estrous stages. Taking a view on the consecutive 16 days of vaginal smears images of TG2, the cell type of day 13 displayed a predominance of cornified enucleate epithelial cells, which was similar to WT representative cell type of estrous stage. But the cell types of day 14 and 15 were similar to the diestrus or proestrus stage due to the predominance of parabasal cells and few cornified superficial cells. Scale bar= 100 μm.

**Fig. S4.**
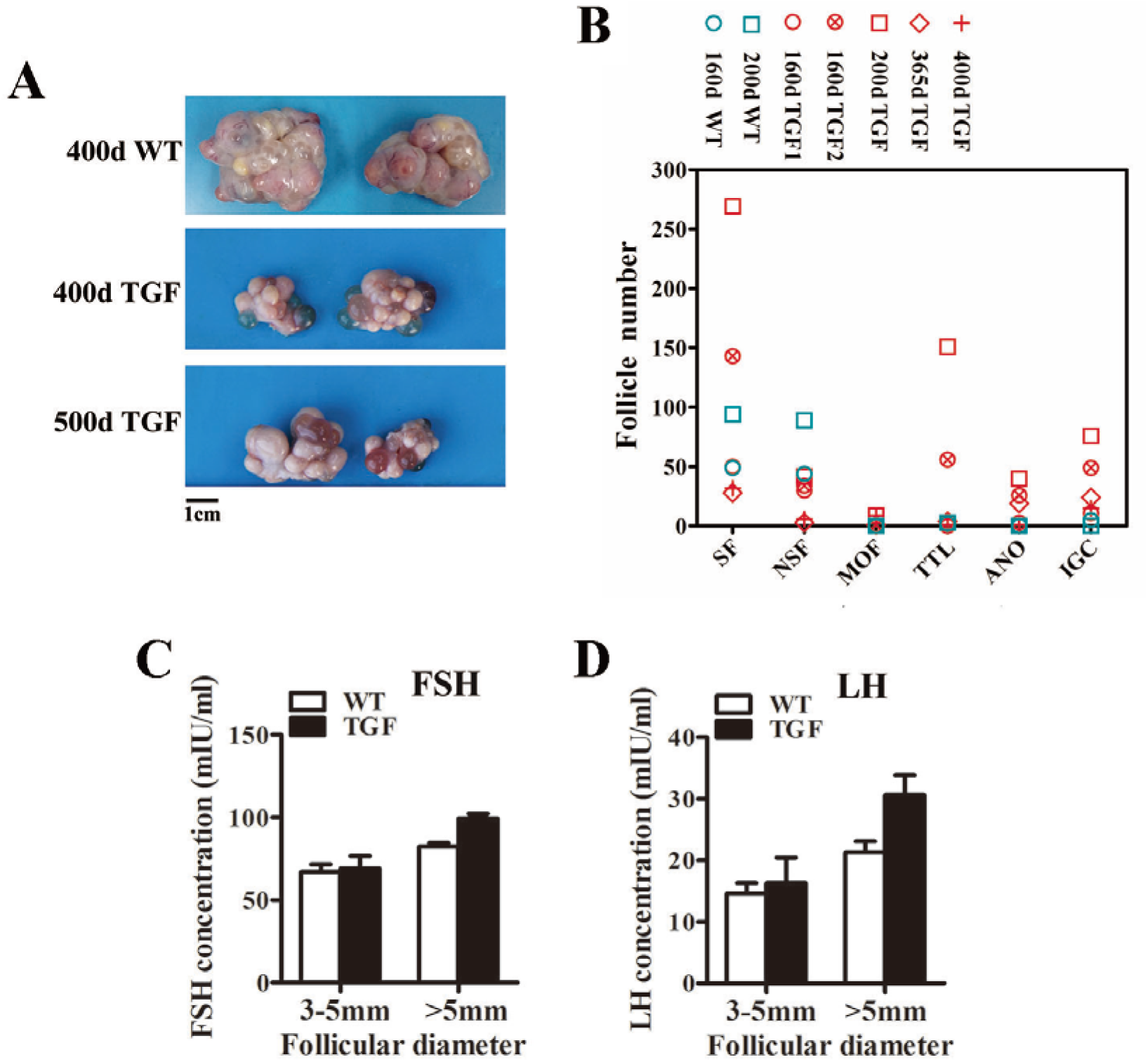
The phenotypes of TGF ovaries. **(A)** Both 400 and 500-day TGF ovaries were apparently smaller than 400-day WT ovaries, but they contained plenty of corpus luteums on the surface. **(B)** Statistical analysis showed less normal secondary follicles (SFs) in TGF ovaries but higher proportion of abnormal SFs. Each three ovarian sections of two WT ovaries and five TGF ovaries were examined. These ovaries were from different gilts at age of 160 to 400 days. Four types of abnormal follicular were distinguished as followings: MOF, multioocyte follicle; TTL, thickened theca and basal lamina; ANO, abnormal oocyte; IGC, irregular and degrading granular cells. NSF stands for normal secondary follicle. **(C)** The concentration of both FSH and LH **(D)** in follicular fluid was not significantly different e between TGF and WT antral follicles.

**Fig. S5.**
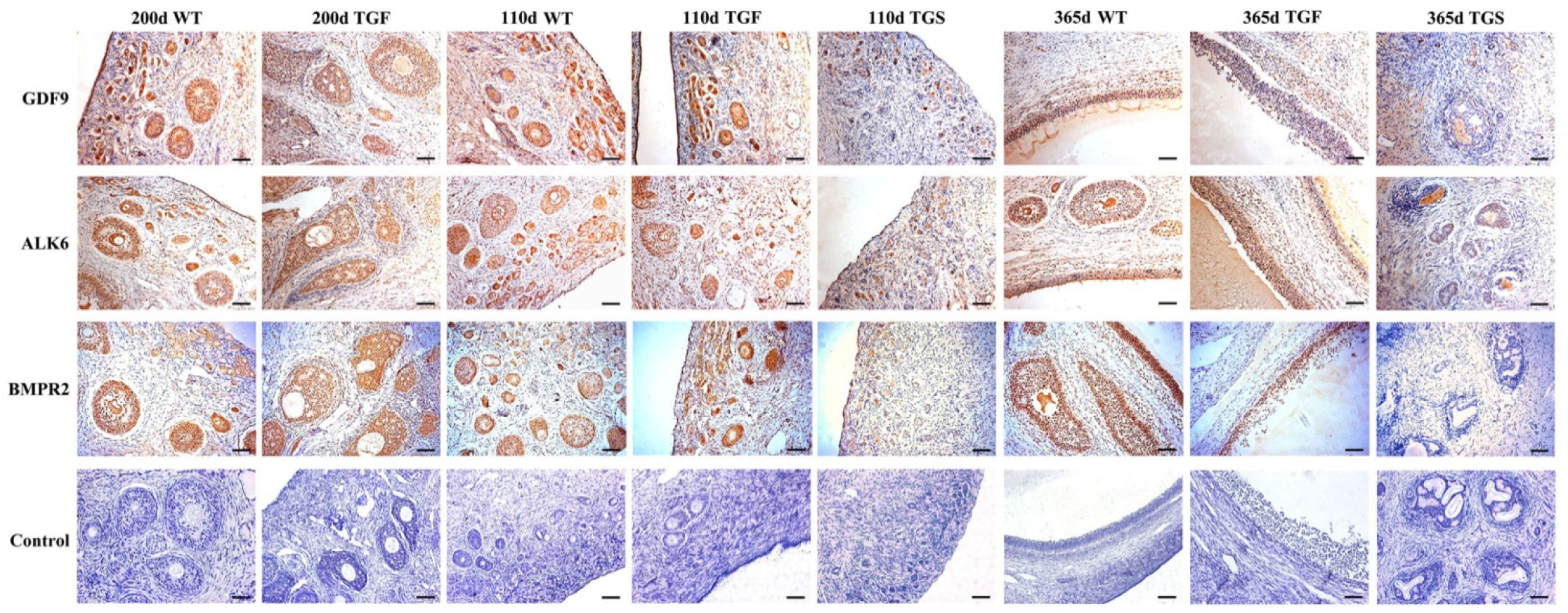
Expression of GDF9, ALK6 and BMPR2 was not affected in TGF follicles. Immunohistochemical staining showed that expression levels of GDF9, ALK6 and BMPR2 were not significant different between TGF and WT follicles, but significantly declined in 110 and 365-day TGS follicles. Scale bar = 100μm.

**Fig. S6.**
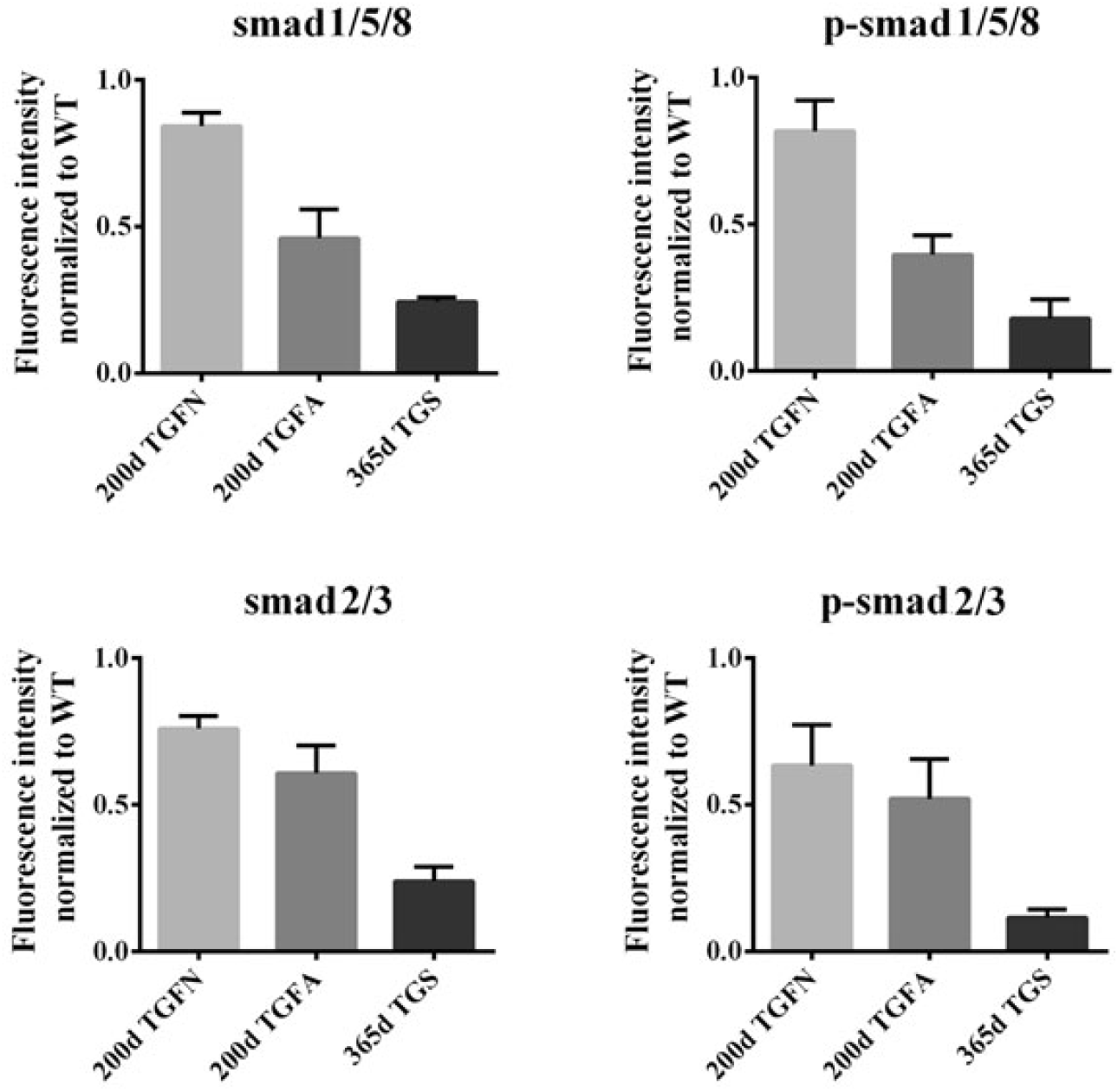
Smad1/5/8 signaling transduction was more affected by knocking down of BMP15 in TGF abnormal follicles. Fluorescence intensity of Smad1/5/8 signaling was decreased more than Smad2/3 signaling in TGF abnormal follicles, though they were both remarkably decreased in TGS follicles. Fluorescence signals were quantied by Image J software, statistic analysis was used graphpad prism software.

**Fig. S8.**
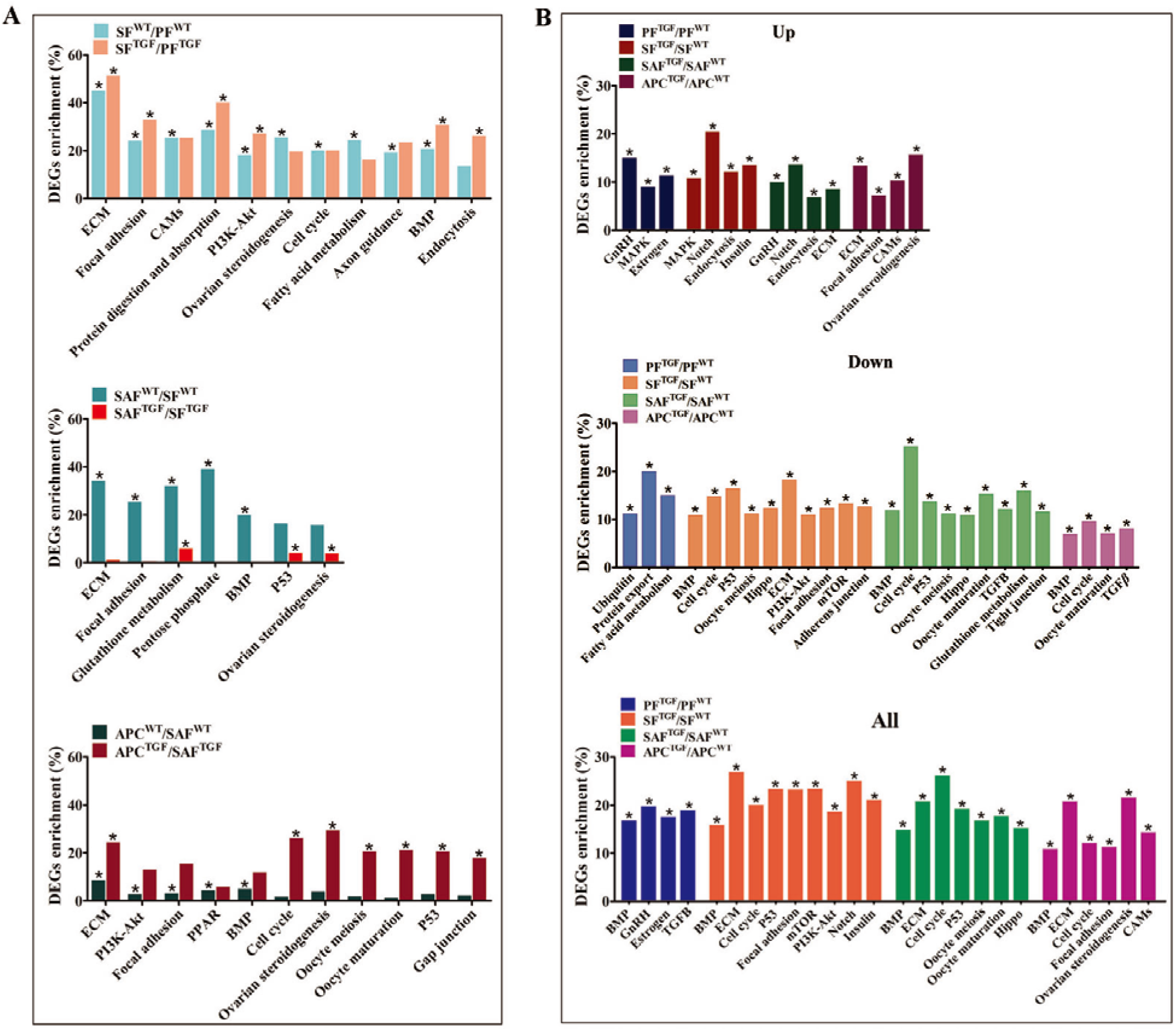
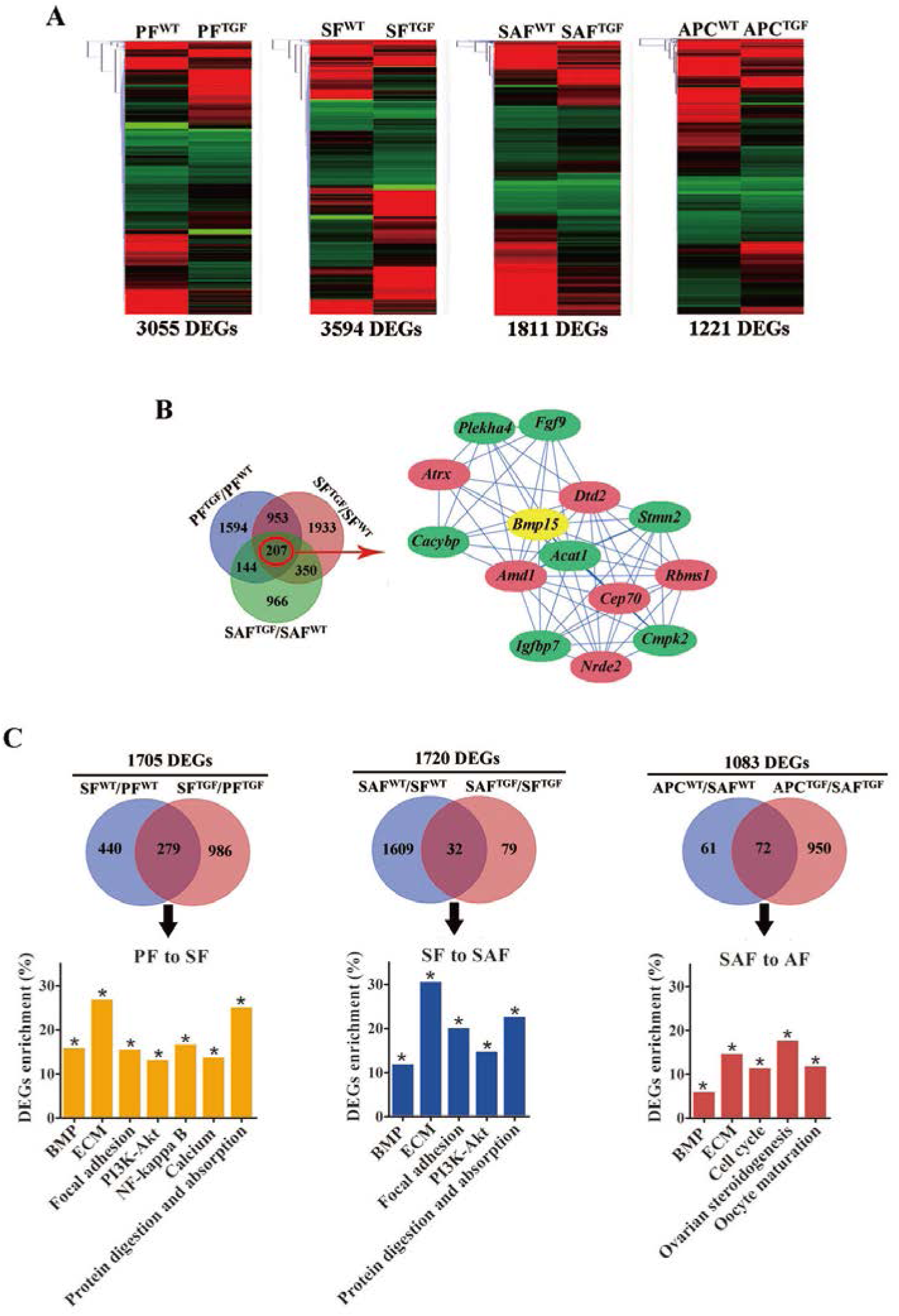
Further analysis of the transcriptomic data. **(A)** Both DEGs number and their clustering pattern revealed a highly different gene expression between WT and TGF follicle during each follicle stages. **(B)** A correlation analysis of the DEGs predicted 7 DEGs (in green ellipse) closely negatively correlated to *Bmp15* (correlation coefficient <-0.98)，and 6 DEGs (in red ellipse) closely positive correlated to *Bmp15* (correlation coefficient >0.98). **c** Further analysis of the three developmental transitions based on twice DEGs identifications.

**Table S1.**
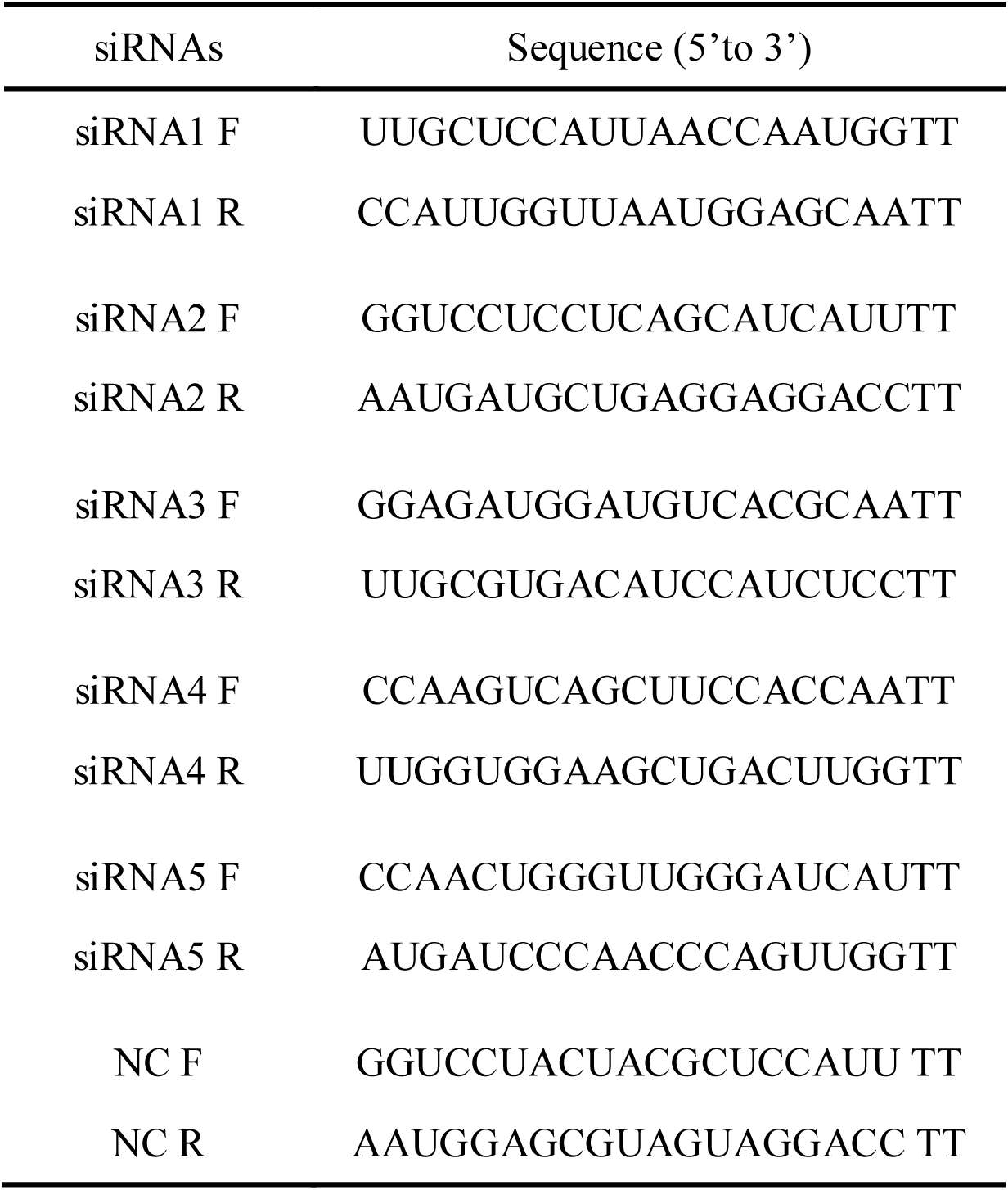
Sequence of siRNA targeting porcine *Bmp15* gene.

**Table S2.**
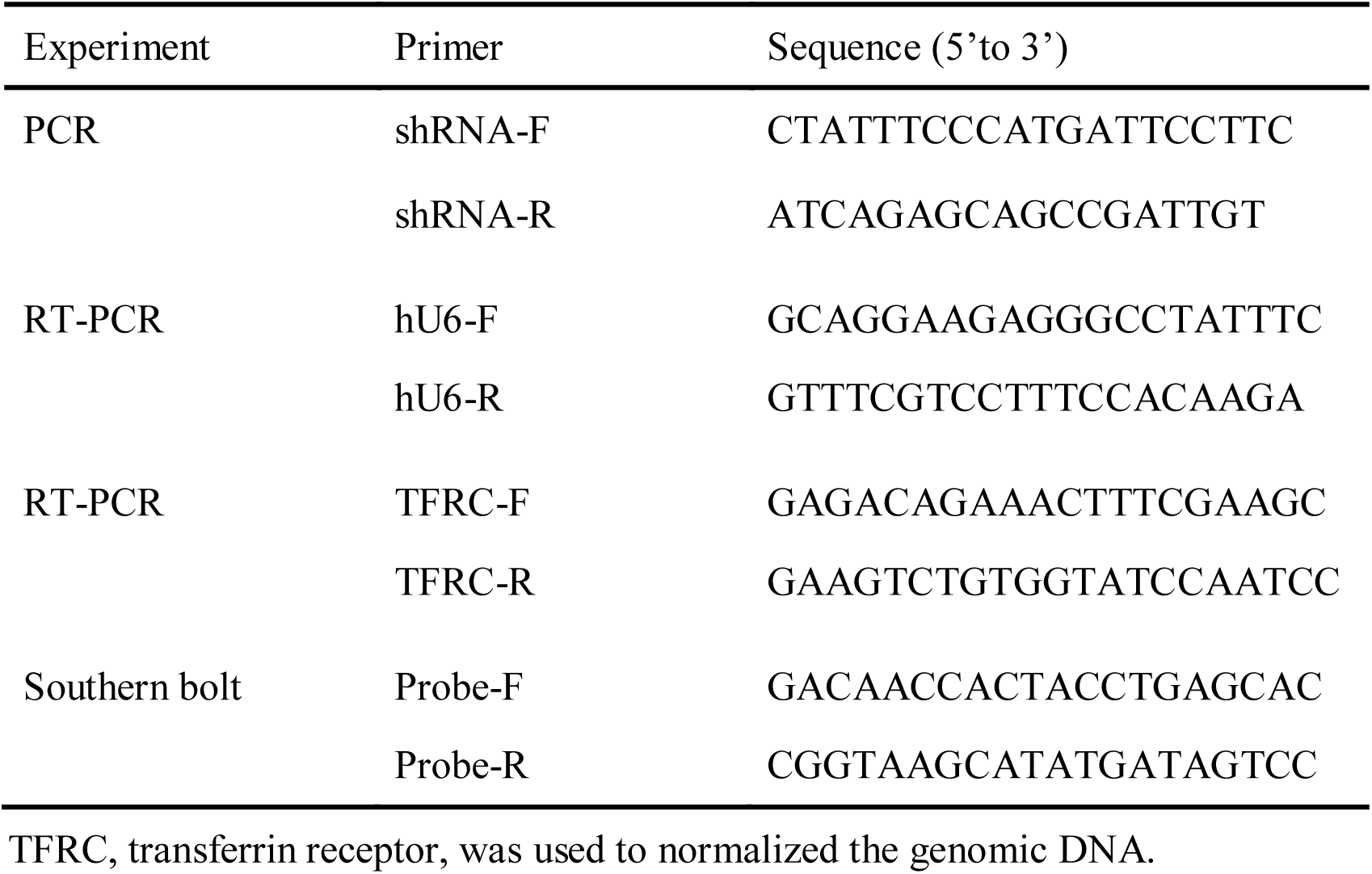
Primers for integrated plasmid detection.

**Table S3.**
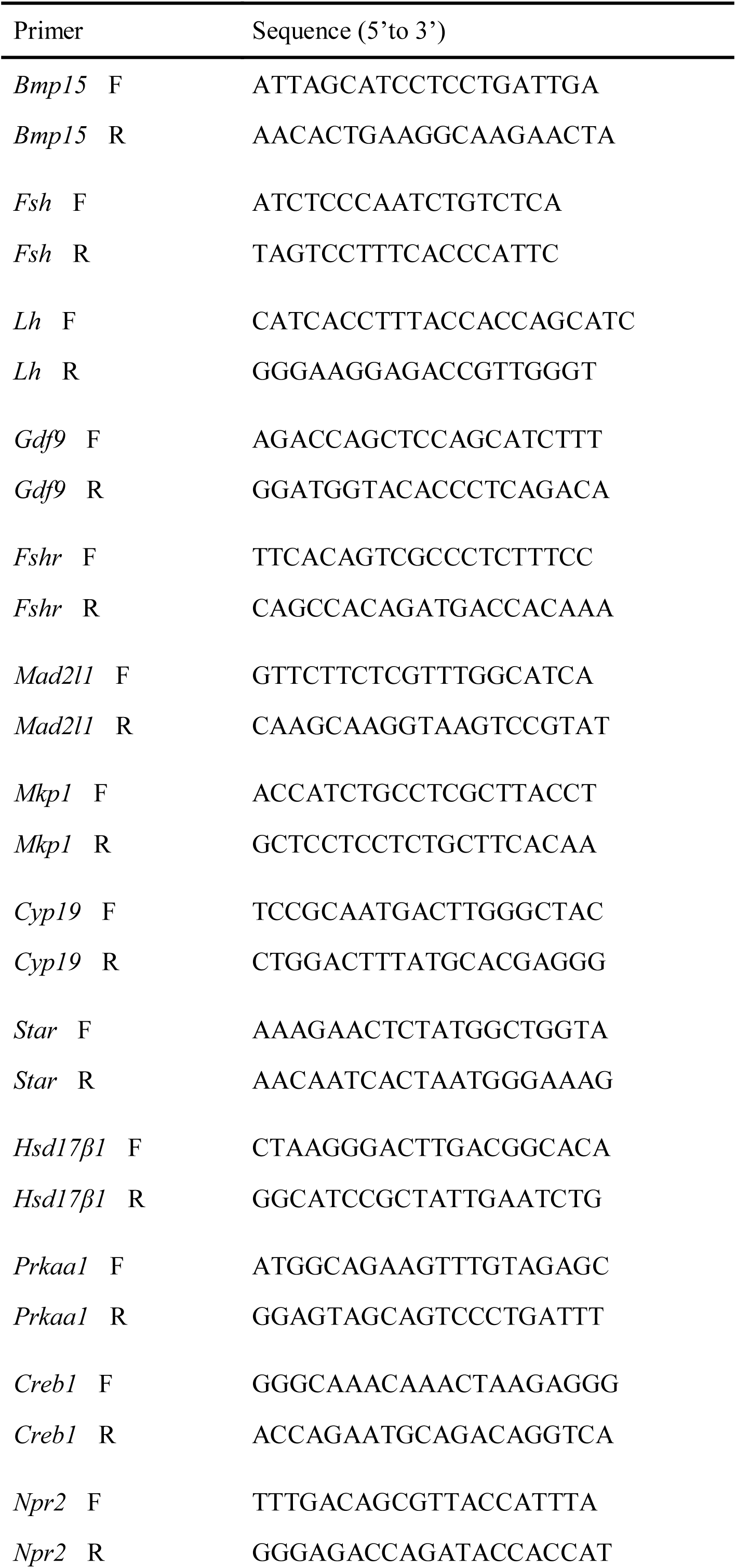
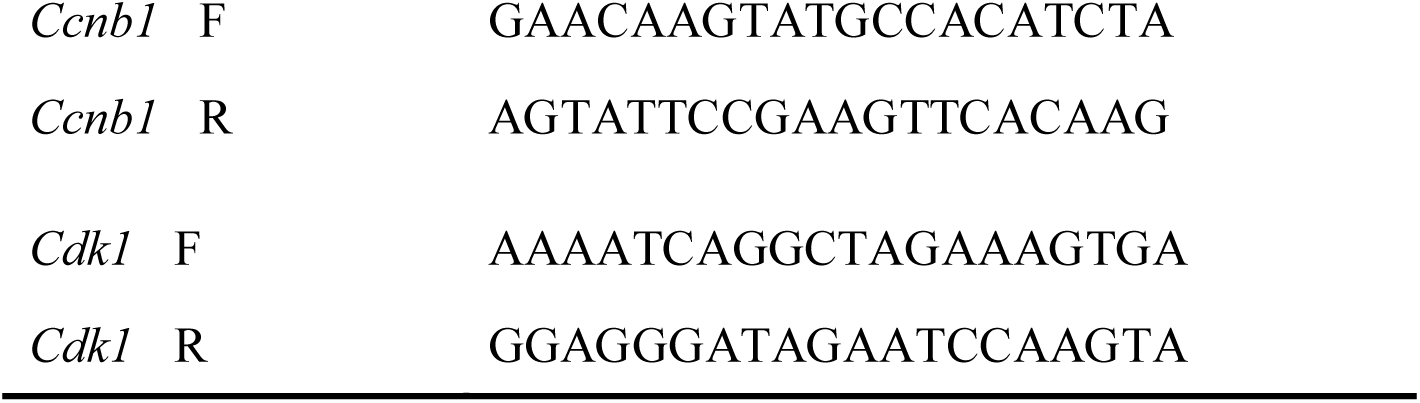
Primers for qPCR detection.

**Table S4.** Weight of ovaries from gilts of different ages.

| Day | Gilt | Weight L (g) | Weight R (g) | Total weight (g) |
| --- | --- | --- | --- | --- |
| 110 | TG | 0.11 | 0.12 | 0.23 |
|  | WT | 0.82 | 0.84 | 1.66 |
| 140 | TG | 0.17 | 0.25 | 0.42 |
|  | WT | 0.55 | 0.72 | 1.27 |
| 150 | TG | 0.16 | 0.13 | 0.29 |
|  | WT | 2.41 | 2.09 | 4.5 |
| 170 | TG | 0.23 | 0.27 | 0.5 |
|  | WT | 1.0 | 0.9 | 1.9 |

L, left ovary ; R, right ovary

**Table S5.**
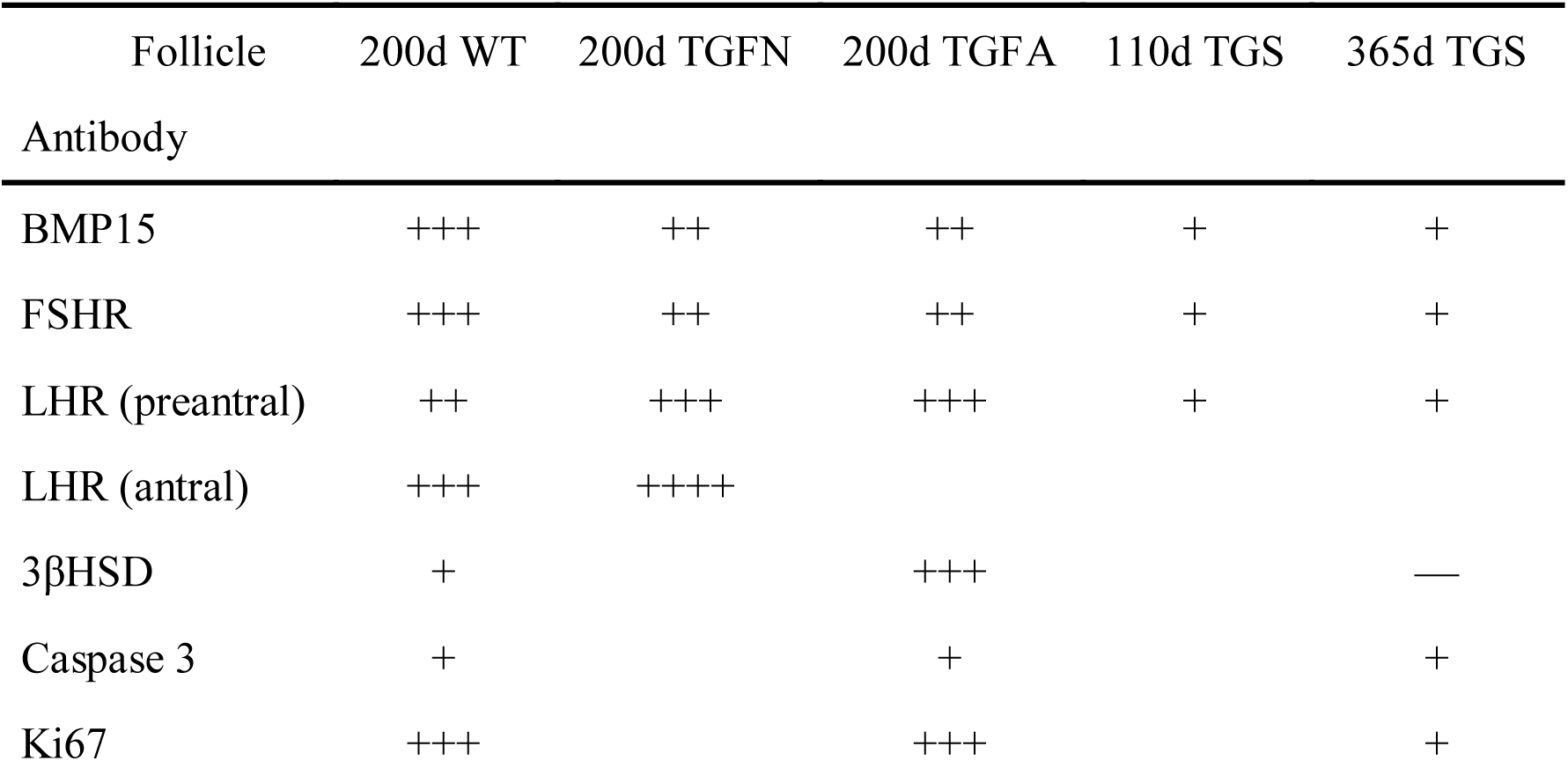
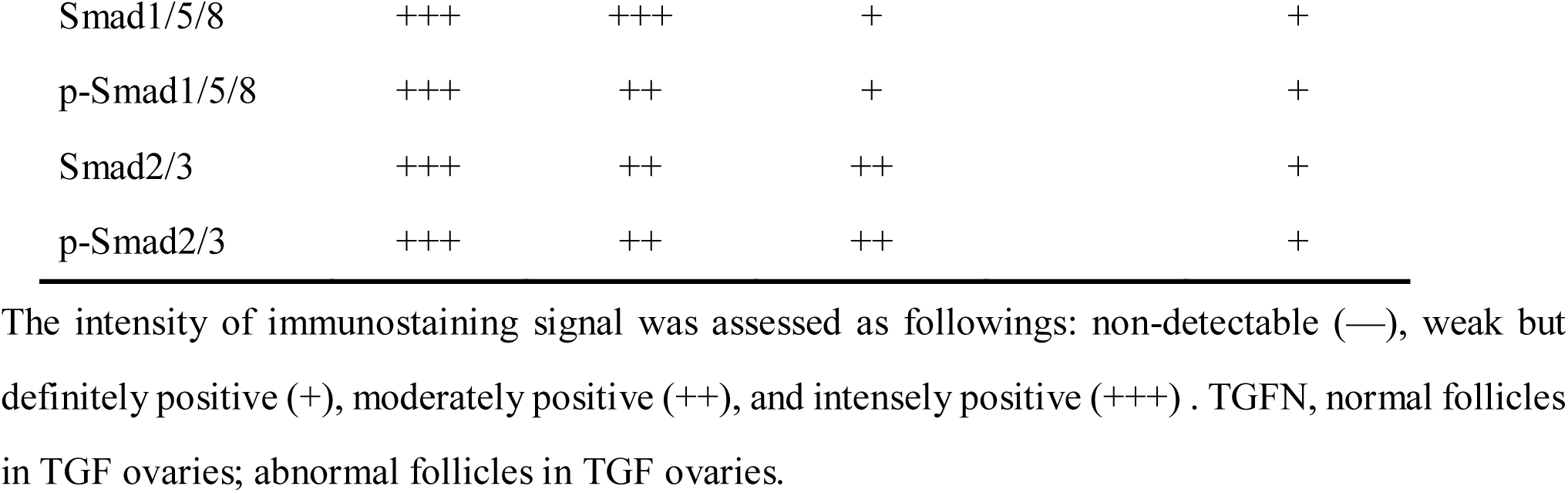
Statistical analysis of intensity of immunostaining signal for each detected factor.

**Table S6.** Summary of DEGs in comparisons.

| Comparisons | Total | Up-regulated | Down-regulated |
| --- | --- | --- | --- |
| Intra effect |  |  |  |
| SF<sup>WT</sup>/PF<sup>WT</sup> | 2877 | 1099 | 1778 |
| SAF<sup>WT</sup>/SF<sup>WT</sup> | 3503 | 1168 | 2335 |
| APC<sup>WT</sup>/SAF<sup>WT</sup> | 350 | 163 | 187 |
| SF<sup>TGF</sup>/PF<sup>TGF</sup> | 4503 | 1594 | 2909 |
| SAF<sup>TGF</sup>/SF<sup>TGF</sup> | 236 | 74 | 162 |
| APC<sup>TGF</sup>/SAF<sup>TGF</sup> | 2390 | 1126 | 1264 |
| Inter effect |  |  |  |
| PF<sup>TGF</sup>/PF<sup>WT</sup> | 3055 | 1482 | 1573 |
| SF<sup>TGF</sup>/SF<sup>WT</sup> | 3594 | 1658 | 1936 |
| SAF<sup>TGF</sup>/SAF<sup>WT</sup> | 1812 | 914 | 898 |
| APC<sup>TGF</sup>/APC<sup>WT</sup> | 1221 | 606 | 615 |

PF, primary follicle; SF, secondary follicle; SAF, small antrum follicle; APC, large antrum parietal cell.

**Table S7.**
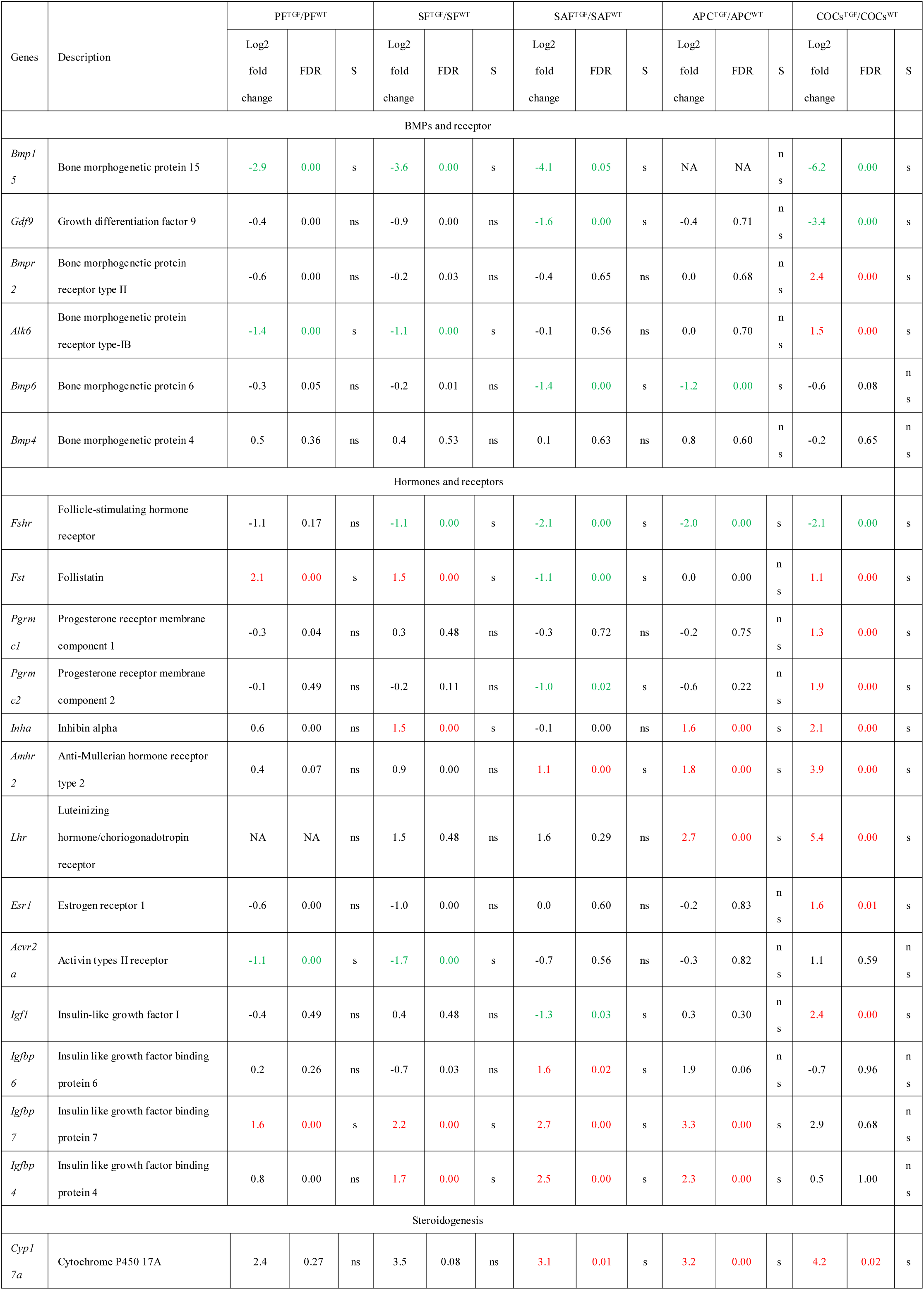
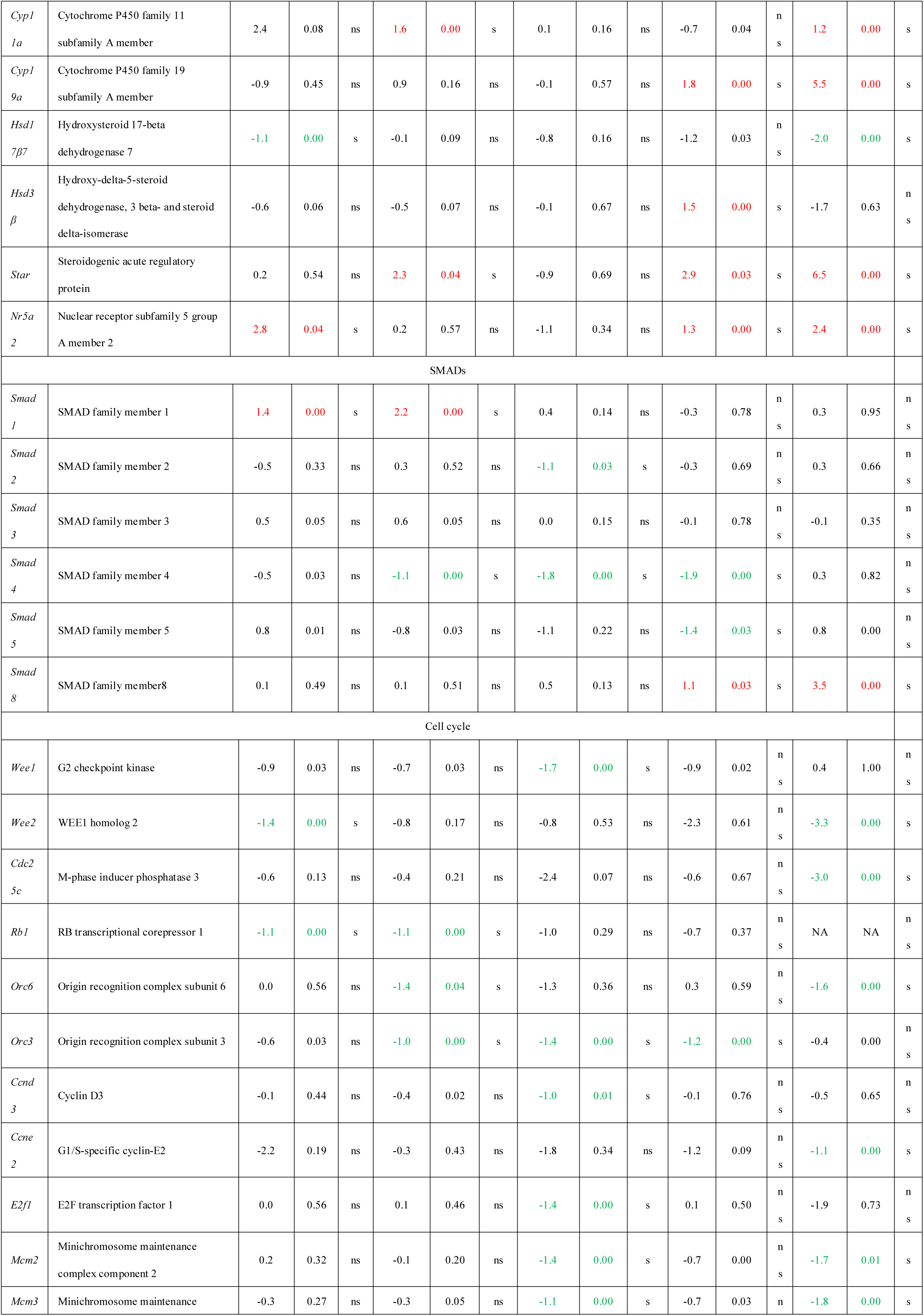
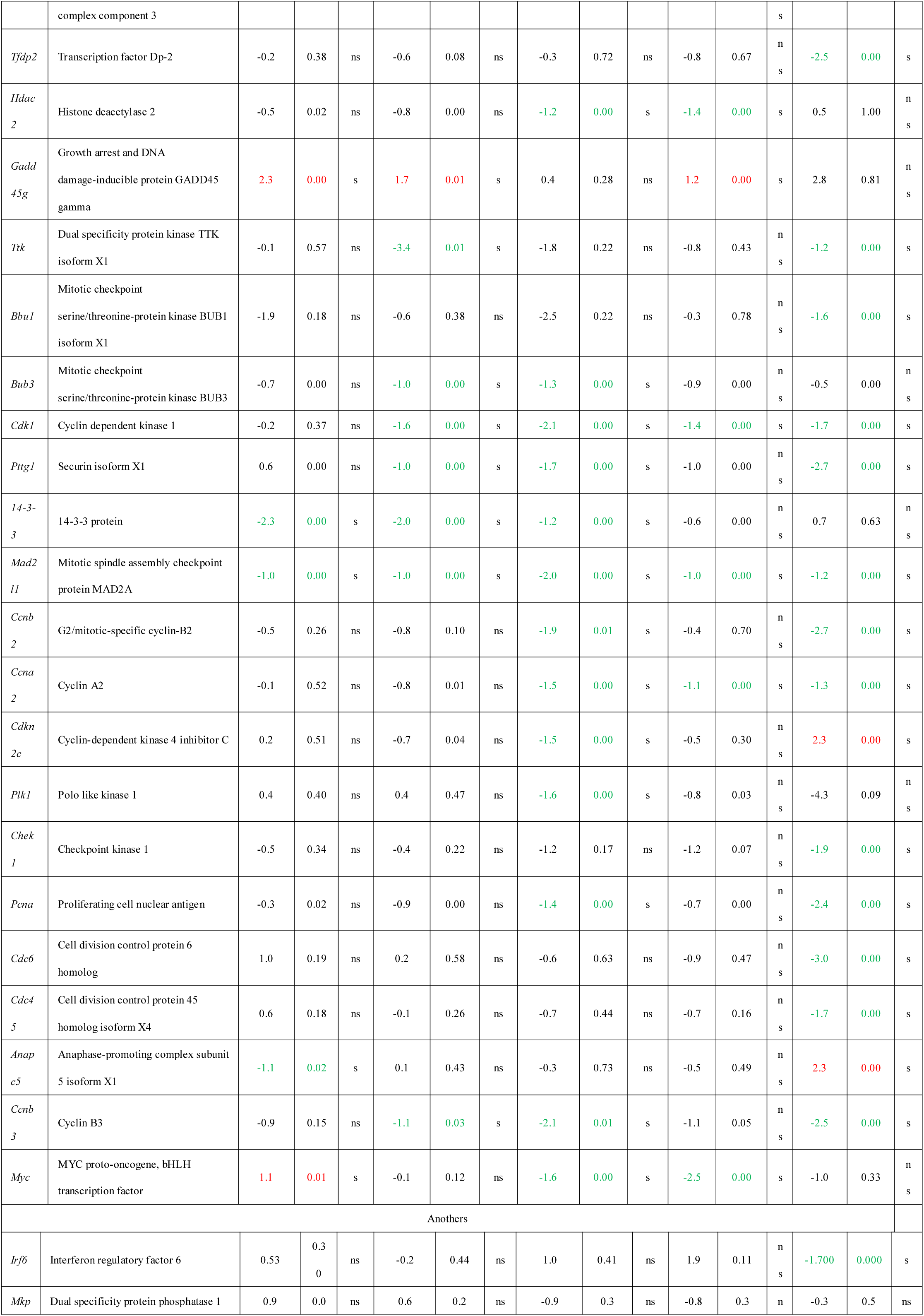
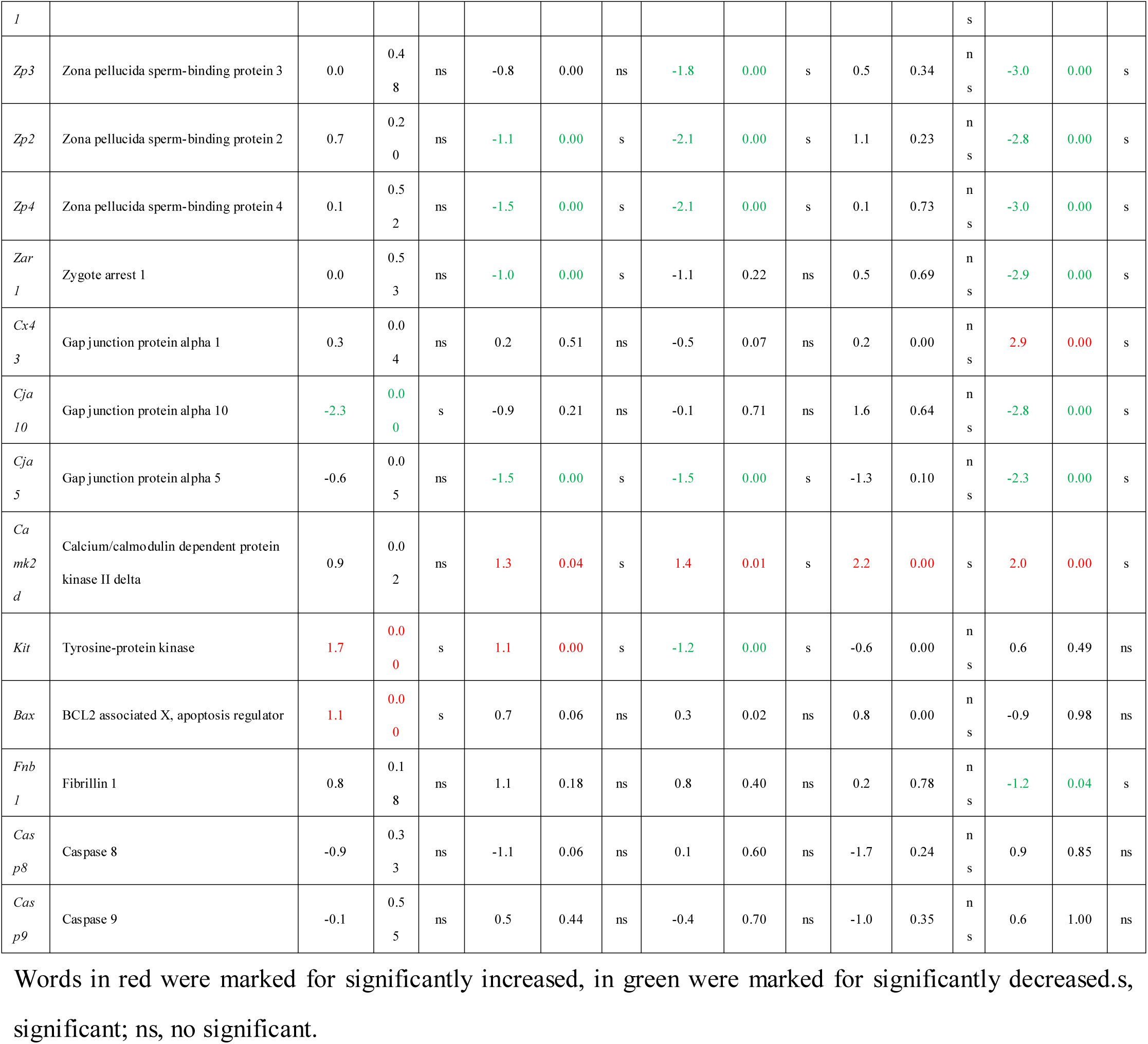
Transcription profile of related genes.

**Table S8.**
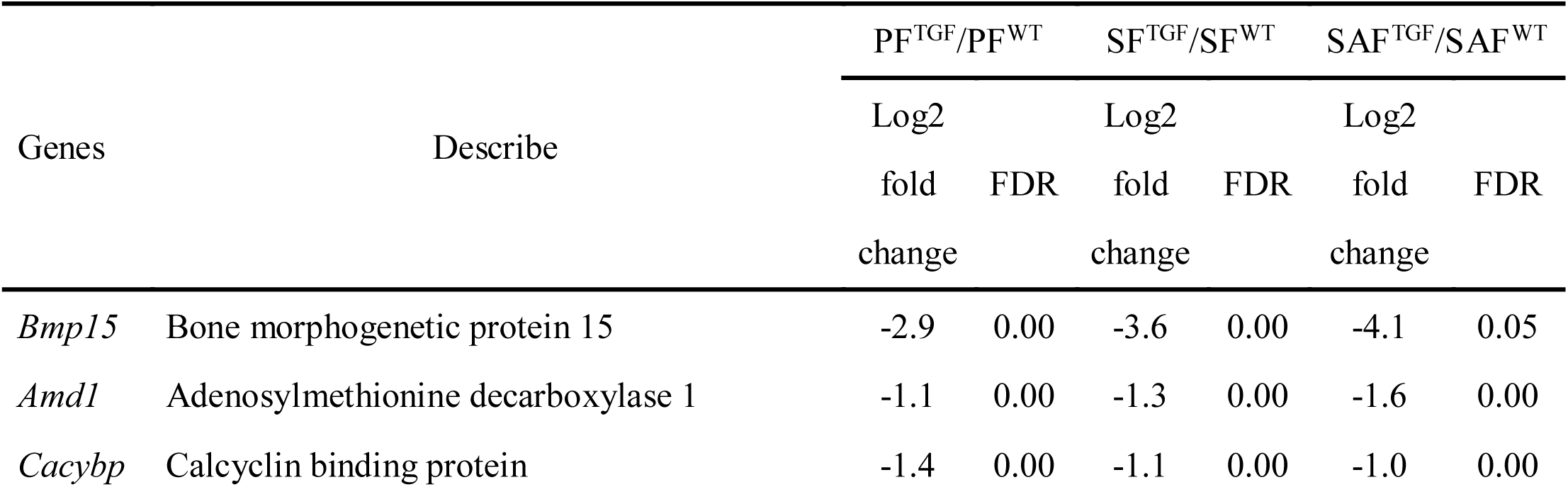
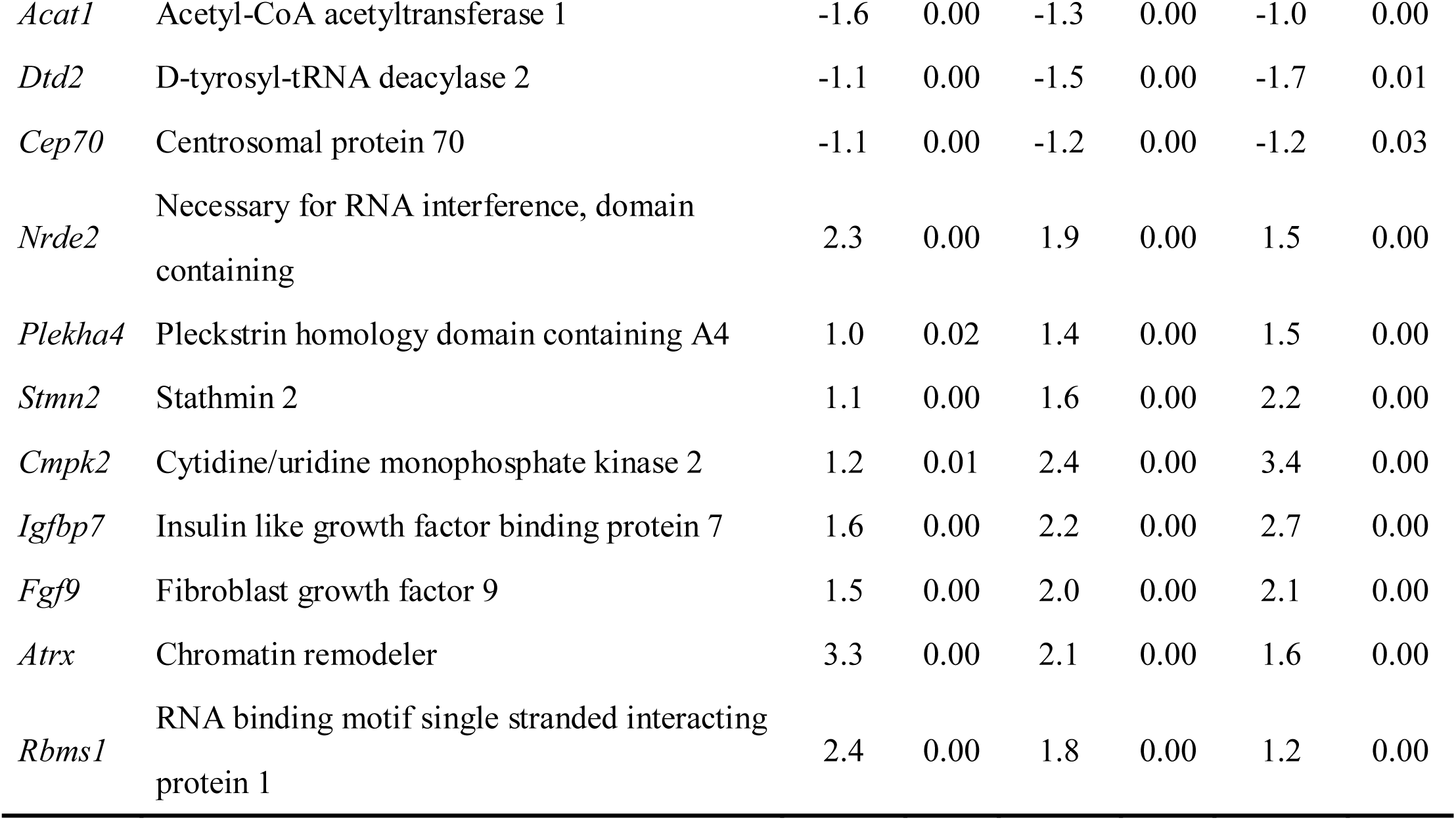
Transcription profile of genes closely correlation with *Bmp15*.

| Genes | Describe | PF<sup>TGF</sup>/PF<sup>WT</sup> |  | SF<sup>TGF</sup>/SF<sup>WT</sup> |  | SAF<sup>TGF</sup>/SAF<sup>WT</sup> |  |
| --- | --- | --- | --- | --- | --- | --- | --- |
|  |  | Log2 |  | Log2 |  | Log2 |  |
|  |  | fold | FDR | fold | FDR | fold | FDR |
|  |  | change |  | change |  | change |  |
| *Bmp15* | Bone morphogenetic protein 15 | -2.9 | 0.00 | -3.6 | 0.00 | -4.1 | 0.05 |
| *Amd1* | Adenosylmethionine decarboxylase 1 | -1.1 | 0.00 | -1.3 | 0.00 | -1.6 | 0.00 |
| *Cacybp* | Calcyclin binding protein | -1.4 | 0.00 | -1.1 | 0.00 | -1.0 | 0.00 |
| *Acat1* | Acetyl-CoA acetyltransferase 1 | -1.6 | 0.00 | -1.3 | 0.00 | -1.0 | 0.00 |
| *Dtd2* | D-tyrosyl-tRNA deacylase 2 | -1.1 | 0.00 | -1.5 | 0.00 | -1.7 | 0.01 |
| *Cep70* | Centrosomal protein 70 | -1.1 | 0.00 | -1.2 | 0.00 | -1.2 | 0.03 |
| *Nrde2* | Necessary for RNA interference, domain containing | 2.3 | 0.00 | 1.9 | 0.00 | 1.5 | 0.00 |
| *Plekha4* | Pleckstrin homology domain containing A4 | 1.0 | 0.02 | 1.4 | 0.00 | 1.5 | 0.00 |
| *Stmn2* | Stathmin 2 | 1.1 | 0.00 | 1.6 | 0.00 | 2.2 | 0.00 |
| *Cmpk2* | Cytidine/uridine monophosphate kinase 2 | 1.2 | 0.01 | 2.4 | 0.00 | 3.4 | 0.00 |
| *Igfbp7* | Insulin like growth factor binding protein 7 | 1.6 | 0.00 | 2.2 | 0.00 | 2.7 | 0.00 |
| *Fgf9* | Fibroblast growth factor 9 | 1.5 | 0.00 | 2.0 | 0.00 | 2.1 | 0.00 |
| *Atrx* | Chromatin remodeler | 3.3 | 0.00 | 2.1 | 0.00 | 1.6 | 0.00 |
| *Rbms1* | RNA binding motif single stranded interacting protein 1 | 2.4 | 0.00 | 1.8 | 0.00 | 1.2 | 0.00 |

**Table S9.**
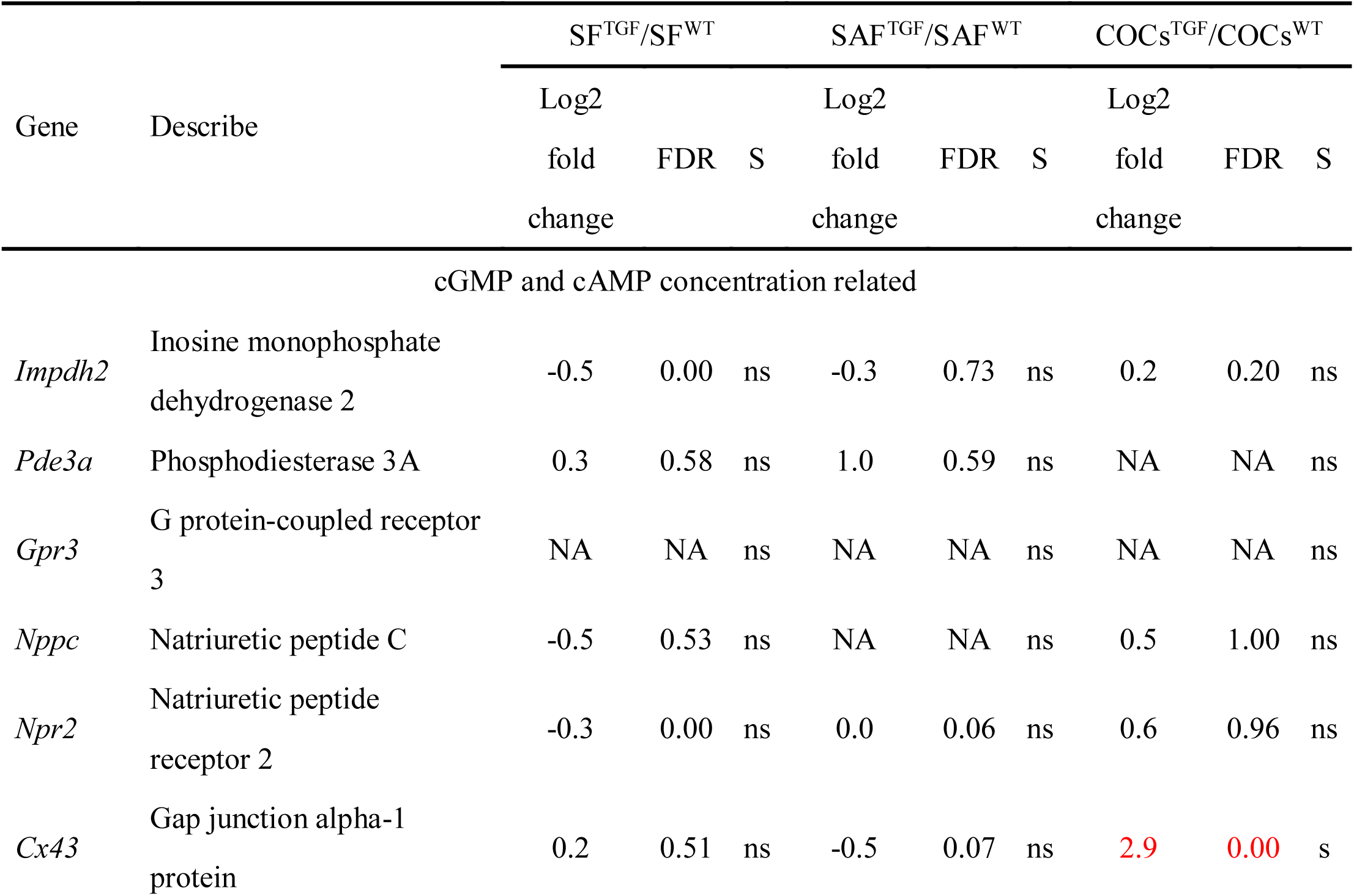
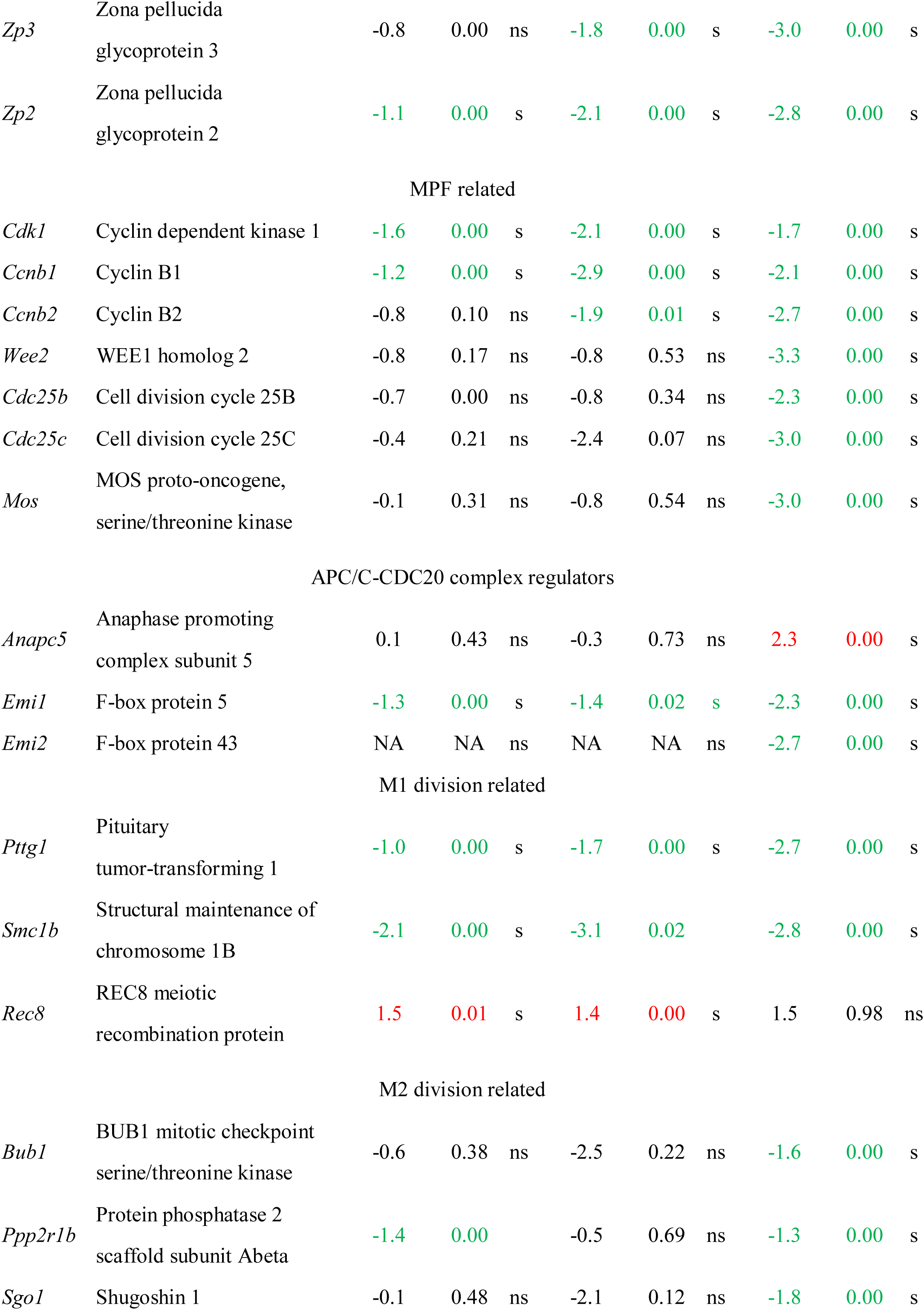
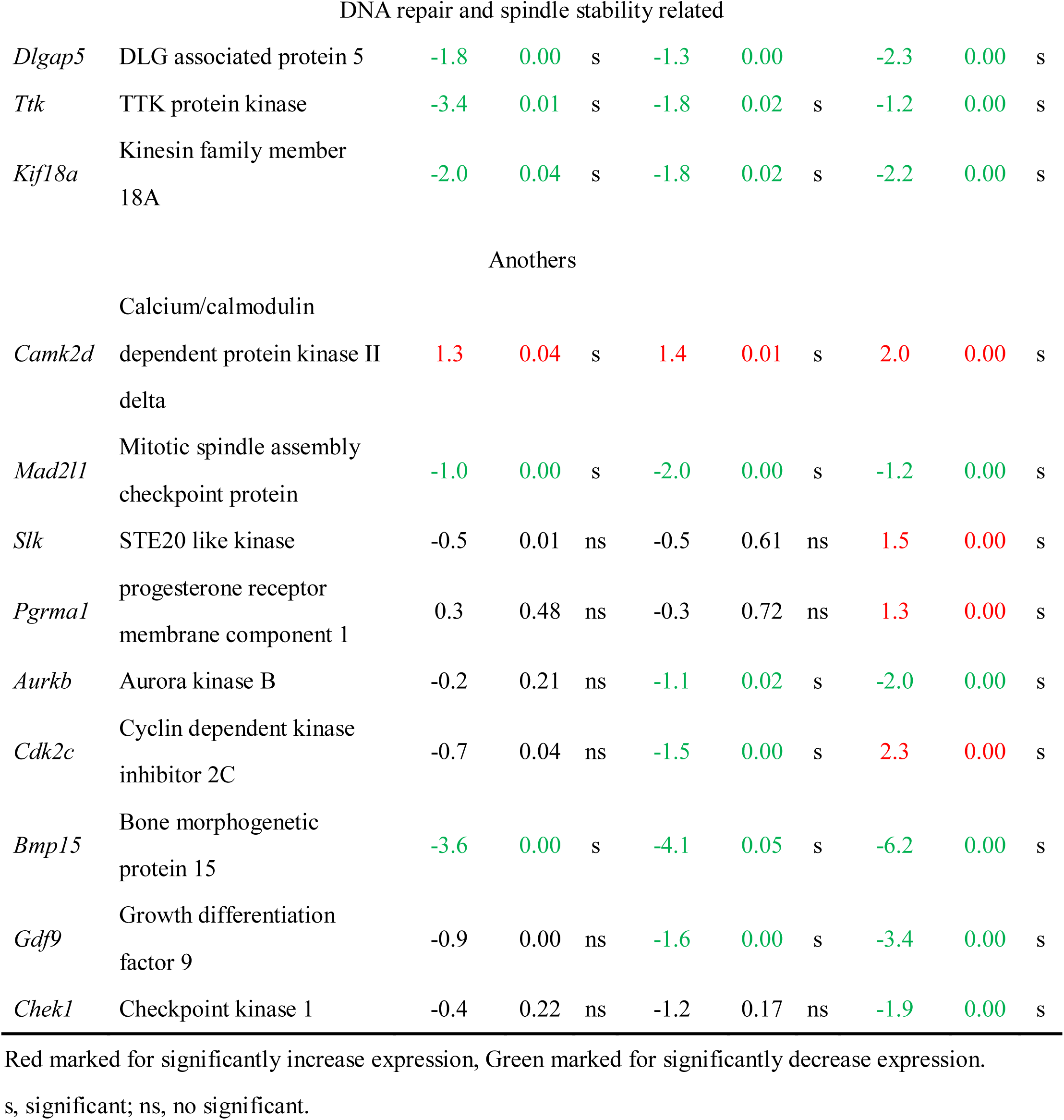
Genes involved in oocyte meiosis.

